# Phase separation-mediated formation of condensed GIT/PIX enzyme complex module for compartmentalized signaling

**DOI:** 10.1101/2019.12.18.881771

**Authors:** Jinwei Zhu, Qingqing Zhou, Yitian Xia, Lin Lin, Jianchao Li, Mengjuan Peng, Rongguang Zhang, Mingjie Zhang

**Affiliations:** State Key Laboratory of Molecular Biology, CAS Center for Excellence in Molecular Cell Science, Shanghai Institute of Biochemistry and Cell Biology, Chinese Academy of Sciences; Shanghai Science Research Center, 333 Haike Road, Shanghai 201210, China; Bio-X Institutes, Key Laboratory for the Genetics of Developmental and Neuropsychiatric Disorders, Ministry of Education, Shanghai Jiao Tong University, Shanghai 200240, China; Division of Life Science, State Key Laboratory of Molecular Neuroscience, Hong Kong University of Science and Technology, Clear Water Bay, Kowloon, Hong Kong, China; School of Life Science and Technology, ShanghaiTech University, 100 Haike Road, Shanghai 201210, China; Center of Systems Biology and Human Health, Hong Kong University of Science and Technology, Clear Water Bay, Kowloon, Hong Kong, China

## Abstract

Cells compartmentalize enzymes for broad physiological functions such as efficient metabolic reactions and spatiotemporally controlled signaling. A given enzyme or enzyme complex can participate in multiple cellular processes in response to different signal inputs by forming different cellular compartments. Here, we demonstrate that association of GIT1 and β-Pix, a pair of GTPase regulatory enzymes involved in diverse cellular processes, leads to autonomous condensation of the complex via phase separation without additional scaffolding molecules. The atomic structure of the GIT/PIX complex reveals the molecular basis governing the phase separation-mediated condensation of the GIT1/β-Pix complex. Importantly, the GIT1/β-Pix condensates can function as a versatile modular membrane-less organelle-like structure for distinct cellular compartmentalization by binding to upstream proteins such as Paxillin in focal adhesions, Shank3 in neuronal synapses, and Scribble in cellular junctions. Thus, phase separation-mediated formation of condensed enzyme complexes provides a powerful way of dynamically concentrating limited amounts of cooperating enzymes to specific cellular compartments for optimal signaling.

## Introduction

In living cells, signals are often initiated, amplified, and transduced at specific subcellular regions with temporal requirements. This requires enzymes to be concentrated at defined subcellular compartments, so that limited amounts of enzymes can satisfy required catalytic activities for proper signaling. Enzymes can be concentrated at specific subcellular regions by binding to their cognate interacting proteins in accordance to traditional thermodynamic binding equilibrium. Emerging evidence suggests that formation of membrane-less compartments, also known as biomolecular condensates, via liquid-liquid phase separation is a new mechanism used by cells to concentrate biomolecules, including enzymes, at specific subcellular regions^1–4^. Membrane-less condensates seem to be widespread in cells and include cellular machineries such as P granule^5^, nucleoli^6^, centrosomes^7,8^, pre- and post- synaptic signaling apparatuses^9–12^, and stress granules^13,14^. Membrane-less biomolecular condensates display many unique features when compared to traditional stoichiometric assemblies of molecular complexes as well as membrane-enclosed cellular compartments^2–4^.

Intuitively, formation of biomolecular condensates can massively enrich both reactants and enzymes within the small volume of a subcellular compartment, and therefore dramatically modify the chemical reactions involved^2,9,15^. Although enormous progress has been made, the mechanisms driving the formation of biomolecular condensates are far from clear. Based on many decades of research on phase separation of chemical polymers and the recent explosive development of the field of biological condensates, one common requirement for forming biomolecular condensates is the presence of multivalent interactions between molecules in each system^2,16,17^. Additionally, it is commonly believed that formation of biomolecular condensates generally require highly abundant organizing molecules such as scaffold proteins, proteins with low complexity sequence, or nucleic acids^2,18^. Although most biological signaling processes are highly specific and sensitive to changes in protein component/concentration under physiological conditions, paradoxically, the majority of reported biological condensates involves or is even dominated by proteins with low complexity/intrinsically disordered sequences that form very weak interactions^19–21^. Several recent studies have provided evidence that highly specific and strong molecular interactions are important for forming various biomolecular condensates such as synaptic signaling machineries^9,10,12^ and cell polarity regulatory complexes^22^.

The GIT and PIX protein families are Arf-specific GTPase-activating proteins (GAPs) and Rho-specific guanine nucleotide exchange factors (GEFs) respectively^23–25^. GIT proteins, including GIT1 and GIT2, share a conserved domain architecture, which consists of an N-terminal zinc finger ArfGAP domain, an ankyrin repeats (ANK) domain, a Spa2-homology domain (SHD), a coiled-coil domain, and a C-terminal focal adhesion (FA) targeting (FAT) domain (Fig. 1a). Each PIX protein, including α-Pix and β-Pix, contains an N-terminal SH3 domain followed by the catalytic Dbl homology (DH) and pleckstrin homology (PH) domain tandem, the GIT-binding domain (GBD), and a C-terminal coiled-coil domain (Fig. 1a). GIT and PIX can each self-associate through their respective coiled-coil domains so that the GIT/PIX complex can form very large molecular mass oligomers^26,27^. Both GIT and PIX can bind to many partner proteins and regulate diverse cellular processes such as FA formation and dynamics, cell polarity and migration, synaptic development and signaling, and immune responses^23^, presumably by functioning as regulatory hubs for the Arf and Rho families of GTPases at specific cellular locations. It is perhaps not surprising that mutations of either *GIT* or *PIX* can cause different human diseases including cancers^28^, psychiatric disorders^29,30^, and autoimmune diseases^31^. Therefore, it is of great importance to understand how the enzyme activities of the GIT/PIX complex might be regulated and how the GIT/PIX complex can modulate diverse cellular processes in different cellular settings.

**Fig. 1.**
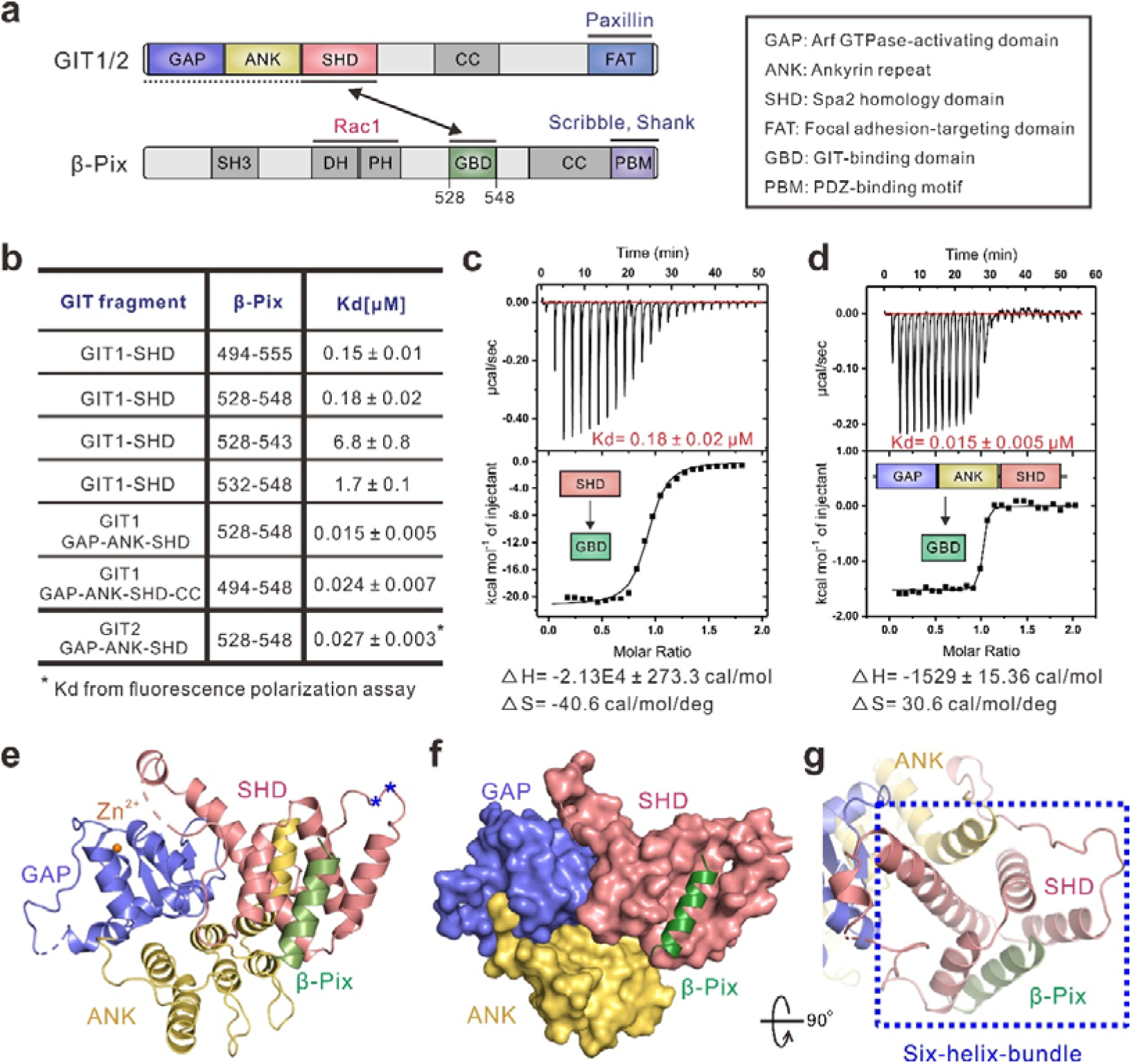
Biochemical and structural characterization of GIT/PIX interaction. (**a**) Schematic diagrams showing the domain organizations of GIT1/2 and β-Pix proteins. The GIT/PIX interaction is indicated by a two-way arrow. The color coding scheme is used throughout the paper. The domain keys are also shown here. (**b**) The dissociation constants of the interactions between various forms of GIT and β-Pix obtained from ITC-based assays. The binding of GIT2 GAP-ANK-SHD (GAS) to β-Pix 528-548 (GBD) was measured using fluorescence polarization assay due to very little heat release of the reaction. (**c,d**) The ITC curves showing the bindings of β-Pix GBD to the isolated SHD domain (**c**) and the GAS tandem (**d**) of GIT1. (**e**) Ribbon diagram representation of the GIT2 GAS/β-Pix GBD complex structure. The residues S255, S256 substituted with Ala during crystal preparation are indicated by blue asterisks. (**f**) The combined surface and ribbon representations of the GIT2 GAS/β-Pix GBD complex structure showing that the GAP, ANK, and SHD domains couple tightly with each other forming a supramodule. (**g**) A six-helix bundle formed by αC^ANK^ of ANK, the four helices from SHD and the β-Pix GBD helix.

In this work, we show that the ArfGAP, ANK, and SHD domains of GIT form an integrated structural supramodule (referred to as the GAS tandem) with a very high binding affinity for the β-Pix GBD (*K*_d_ ∼15 nM). The crystal structure of GIT2 GAS in complex with a synthetic β-Pix GBD peptide solved at the 2.8 Å resolution provides a mechanistic explanation to the tight binding between the two proteins. We further show that the GIT1/β-Pix complex undergoes phase separation, forming highly concentrated enzyme condensates both *in vitro* and in living cells without additional scaffolding molecules. We demonstrate that the GIT1/β-Pix condensates can be recruited to FAs or synapses via binding to Paxillin or Shank3, respectively. Point mutations introduced to GIT1 or β-Pix that specifically perturb the phase separation of the GIT1/β-Pix complex, but not alter the direct interaction between the two enzymes, led to impaired enrichment of the complex in FAs and synapses, and consequent defective migrations of HeLa cells and synapse formation in cultured neurons. Therefore, the GIT/PIX enzymatic complexes can autonomously form modular biomolecular condensates capable of being targeted to specific subcellular compartments by different upstream adaptor proteins to perform distinct cellular functions. The formation of such modular enzymatic condensates can be a mean for cells to concentrate limited amounts of enzymes for performing diverse functions with spatial and temporal features.

## Results

### The GAP-ANK-SHD tandem of GITs binds to β-Pix with a very high affinity

An earlier study reported that the SHD domain of GIT1 binds to a fragment of β-Pix (aa 496-554)^27^. We verified this interaction using purified proteins. Isothermal titration calorimetry (ITC)-based assay showed that the SHD domain of GIT1 binds to β-Pix^494–555^ with a dissociation constant (*K*_d_) of ∼0.15 μM (Fig. 1b). A 21-residue fragment of β-Pix (aa 528-548; referred to as GBD for GIT-binding domain) was found to be sufficient for binding to GIT1 SHD (*K*_d_ ∼0.18 μM) (Fig. 1b and 1c). Further truncation of β-Pix GBD at either end impaired its binding to GIT1 SHD (Fig. 1b), thus the 21-residue GBD is the minimal and complete GIT1 binding region of β-Pix. Unexpectedly, we found that a longer fragment of GIT1, which includes the GAP, ANK, and SHD domains (i.e. the GAS tandem), binds to β-Pix GBD with a ∼10-fold higher affinity than SHD alone does (*K*_d_ ∼0.015 μM; Fig. 1b and 1d), indicating that the GAP, ANK, and SHD domains of GIT1 may form a structural supramodule for binding to β-Pix. We further showed that the GIT2 GAS tandem bound to β-Pix GBD with a similar affinity (*K*_d_ ∼0.027 μM, Fig. 1b).

### Structural basis governing the specific GIT/PIX interactions

To elucidate the molecular basis underlying the specific GIT/PIX interactions, we tried to crystallize β-Pix GBD in complex with the GAS tandem of GIT1 or GIT2. We were able to obtain crystals of the GAS tandem of GIT2 (but no luck with GIT1) in complex with a synthetic β-Pix GBD peptide, but the complex crystals only diffracted to ∼4-5 Å resolution. After numerous trials, we discovered that GIT2 GAS/β-Pix GBD complex prepared from a GIT2 GAS mutant bearing two point mutations in a predicted loop region (S255A/S256A, denoted as GAS^S255A/S256A^; the complex is referred to as the GIT2-GAS/β-Pix GBD complex here on for simplicity) could crystalize and crystals were diffracted to a 2.8 Å resolution. The complex structure was solved by the molecular replacement method using the structure of GAP-ANK tandem of ACAP1 (PDB code: 3JUE) as the searching model (Table S1).

The structure of the GIT2-GAS/β-Pix GBD complex explains how the GAP, ANK, and SHD domains of GIT1 form a supramodule with an enhanced binding to β-Pix GBD. The structure of the GIT1 GAP domain in the complex is very similar to that of the ASAP3 GAP domain observed in a Arf6/ASAP3 complex^32^ (Supplementary Fig. 1a and 1b). Superimposition of the structure of GAP^GIT2^ with that of GAP^ASAP3^ shows that a conserved arginine of GAP^GIT2^, R39^GIT2^, aligns well with the arginine finger of ASAP3, R469^ASAP3^, which is required for GTP hydrolysis (Supplementary Fig. 1b). This structural analysis is consistent with previous findings showing that GIT proteins possess GAP activities toward Arf1 and Arf6^24,33^ and R39 is critical for the GAP activity^34,35^. The ANK domain contains three ankyrin repeats and a C-terminal α-helix (αC^ANK^), which takes a ∼90° bend toward α3B of ANK (Supplementary Fig. 1c). The concave groove of ANK is unoccupied and thus available for potential target binding (Fig. 1e and Supplementary Fig. 1c). The SHD domain in the complex is formed by four consecutive α-helices (α1-α4) (Supplementary Fig. 1d).

In line with our biochemical data, the GAP, ANK, and SHD domains of GIT2 interact with each other intimately to form a structural supramodule (Fig. 1e and 1f). It is noted that S255 and S256 are located at the loop between ANK and SHD and are away from the GIT2/β-Pix interface (highlighted with asterisks in Fig. 1e), so the mutations used to facilitate crystallization should not affect the structure of the GAS tandem and its binding to β-Pix. In the complex, β-Pix GBD forms an α-helix and interacts with GIT2 SHD. A stable six-helix bundle is formed by the β-Pix GBD α-helix, four helices of GIT2 SHD, and αC^ANK^ (Fig. 1g), indicating that the formation of the GITs GAS supramodule may stabilize the conformation of the SHD domain and thus enhance the interactions between GITs and β-Pix.

The GIT2/β-Pix interaction is mainly mediated by hydrophobic interactions. I535, L536, V538, I539 and Y542 from β-Pix GBD form hydrophobic contacts with L270, L273, L277, L281, V285, L334, F337, L345, and I349 from GIT2 SHD (Fig. 2a and 2b). Additional polar interactions (e.g. the hydrogen bond formed between Y542^β-Pix^ and D348^GIT2^, Fig. 2a) further support the binding specificity of the complex. Importantly, the residues involved in the binding interface are highly conserved in both GIT and PIX (Fig. 2c and Supplementary Fig. 2), implying indispensable roles of the GIT/PIX interaction in the animal kingdom. A series of mutations on GIT2 GAS and β-Pix GBD were generated to verify the roles of these residues for the complex formation. Specifically, substitution of L273^GIT2^ or L281^GIT2^ with Ala impaired its binding to β-Pix GBD (Fig. 2d and 2f). Reciprocally, neither the I535D nor the V538D mutants of β-Pix GBD were capable of binding to GIT2 GAS (Fig. 2e and 2f). Substitution of Y542^β-Pix^ to Asp also impaired its binding to GIT2 GAS (Fig. 2e and 2f).

**Fig. 2.**
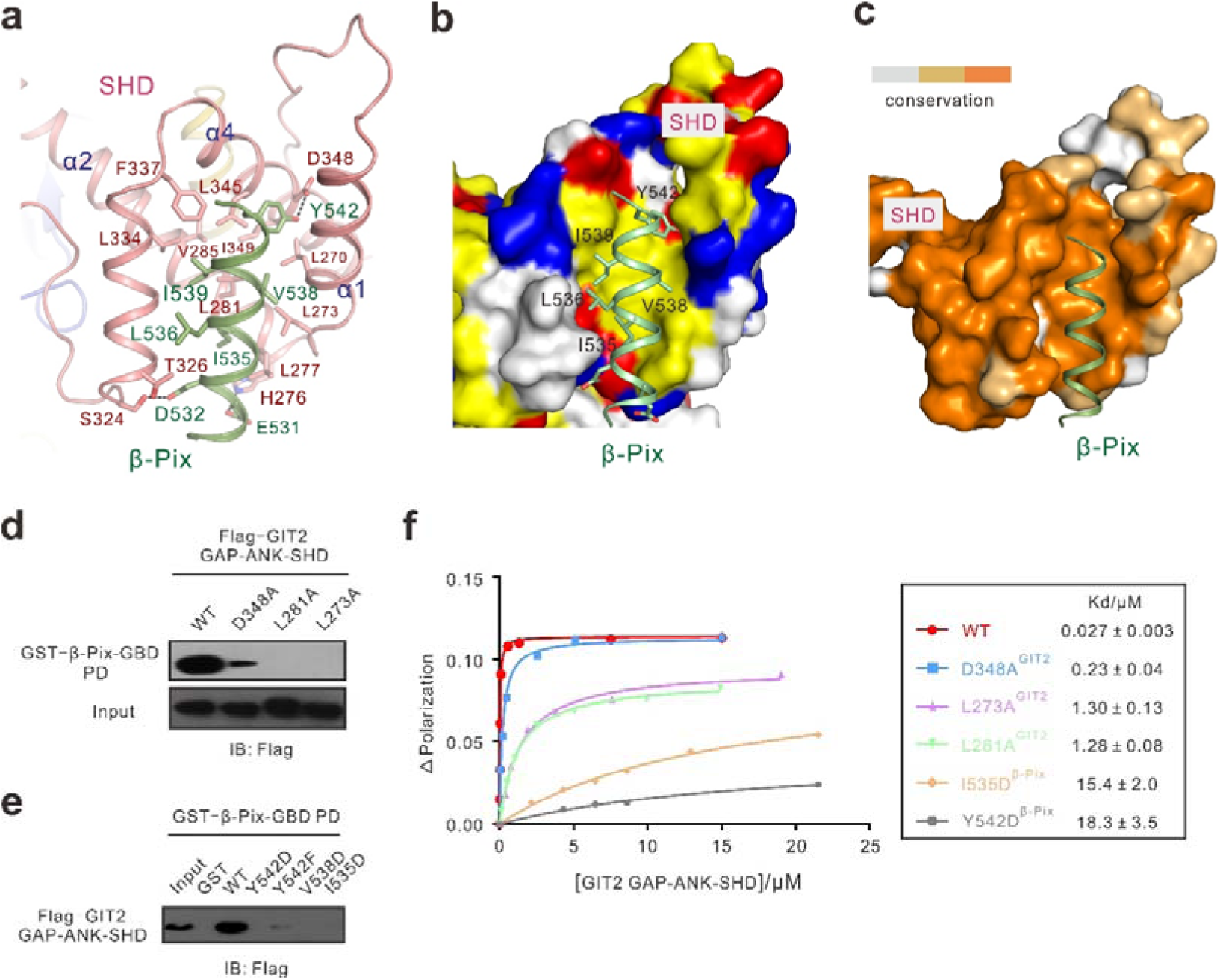
Structural detail of GIT2/β-Pix interface. (**a**) Detailed interactions between GIT2 GAS and β-Pix GBD. (**b**) The combined surface and ribbon representations of the GIT2/β-Pix interface showing that the binding is mainly mediated by hydrophobic interactions and supplemented by hydrogen bonds. In the surface diagram, hydrophobic residues are in yellow, positively charged residues in blue, negatively charged residues in red, and the rest in grey. (**c**) The conservation map of the β-Pix GBD binding site on GIT2 in the surface model showing that the GIT2/β-Pix interaction is highly conserved. (**d,e**) GST-pull down assays showing that key residues involved in the GIT2 GAS/β-Pix GBD interface are required for the interaction. (**f**) Fluorescence polarization-based measurement of binding affinities of WT and mutant GIT2 GAS to WT and mutant β-Pix GBD peptides.

### GIT1 undergoes phase separation in vitro and in cells

Interestingly, we observed that the full-length WT GIT1 solution turned turbid above certain concentrations at room temperature. The turbid solution became clear again upon cooling the protein sample on ice. We sparsely labeled purified GIT1 with Cy3 fluorophore and investigated the turbid solution under a confocal microscope. Cy3-GIT1 formed phase separated droplets with spherical shapes in a concentration dependent manner (Fig. 3a). Differential interference contrast (DIC) microscopic images further showed that small droplets could fuse with each other into larger ones over time (Fig. 3b). Fluorescence recovery after photobleaching (FRAP) analysis of Cy3-GIT1 droplets showed that GIT1 molecules can freely exchange between the condensed and dilute phases (Fig. 3c). Next, we tested whether GIT1 can undergo phase separation in living cells. When GFP-GIT1 was overexpressed in HeLa cells, spherical GFP-GIT1 puncta were observed (Fig. 3d). FRAP assay showed that GFP-GIT1 within these puncta can exchange with the surrounding dilute cytoplasmic population (Fig. 3e), indicating that GFP-GIT1 formed membrane-less condensates in cells.

**Fig. 3.**
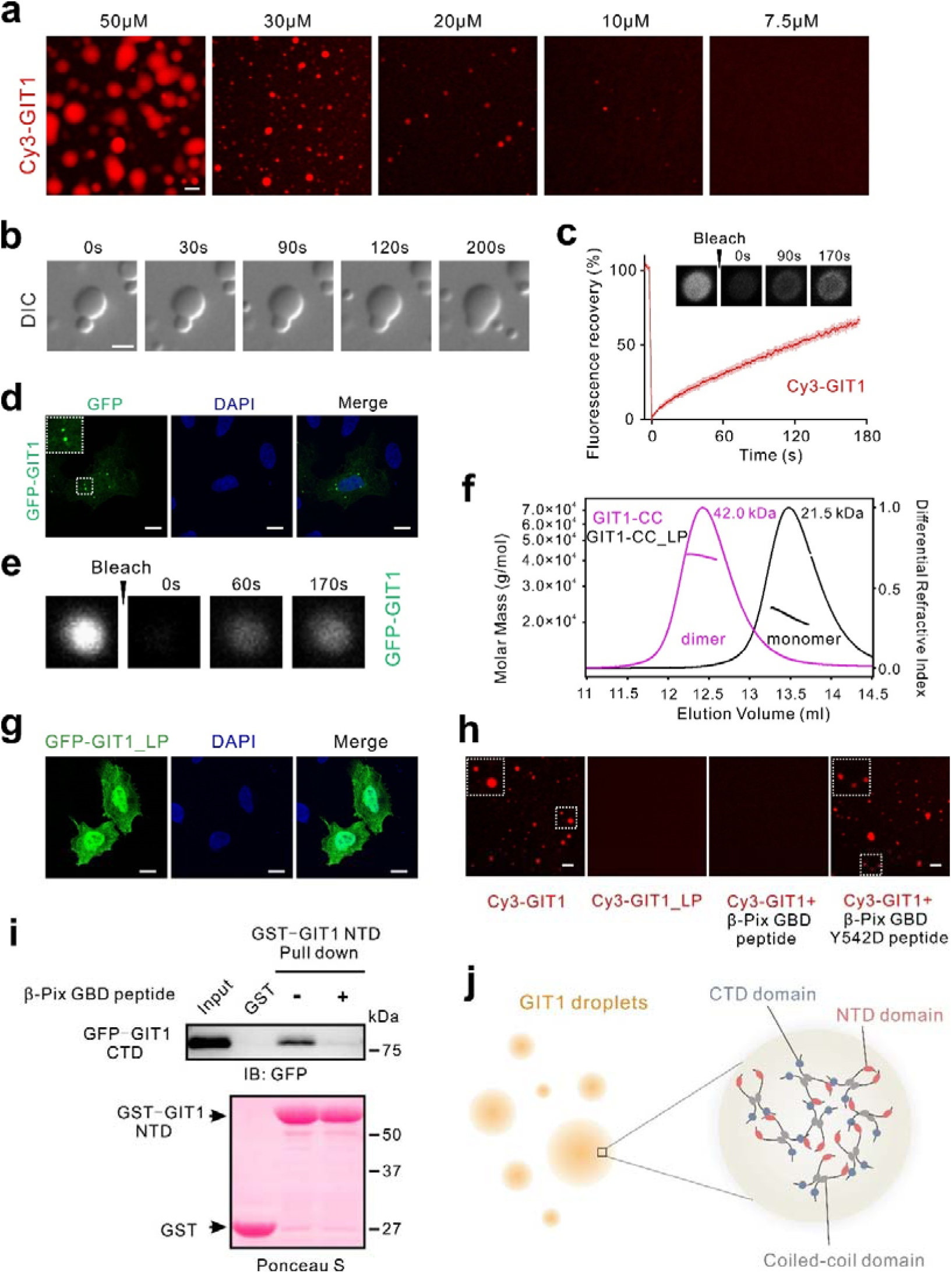
GIT1 undergoes phase separation *in vitro* and in living cells. (**a**) Fluorescence images showing that the full length GIT1 protein underwent phase separation at indicated concentrations. GIT1 was sparsely labeled by Cy3 at 1%. (**b**) DIC images showing that GIT1 condensed droplets fused with each other forming larger droplets over time. (**c**) FRAP analysis showing that GIT1 in the condensed droplets dynamically exchanged with those in the dilute phase. (**d**) Representative images showing expression of GFP-GIT1 in HeLa cells produced many bright and spherical puncta. (**e**) FRAP analysis showing that GFP-GIT1 in the spherical puncta dynamically exchanged with those in cytoplasm. (**f**) FPLC-coupled static light-scattering analysis showing that wild type GIT1-CC formed a stable dimer in solution, whereas the ‘‘LP’’ mutant of GIT1-CC is a monomer. (**g**) Fluorescence images showing that the GIT1_LP mutant cannot form phase separation at ae concentration of 20 μM. Binding of β-Pix GBD peptide, but not β-Pix GBD_Y542D peptide, abolished the phase separation of GIT1. (**h**) Representative images showing expression of GFP-GIT1_LP mutant in HeLa cells did not form any puncta. (**i**) GST-pull down assay showing that the N-terminal fragment (aa 1-371, NTD) of GIT1 binds to the C-terminal fragment of GIT1 (aa 371-end, CTD). Addition of β-Pix GBD peptide impaired the interaction between NTD and CTD of GIT1. (**j**) A model depicting the mechanism of GIT1 phase separation. Scale bar: 5 μm.

We next dissected the molecular mechanism that governs GIT1 phase separation. In many biological systems, proteins containing intrinsic disordered regions (IDRs) can phase separate under physiological conditions^2,3,36^. GIT1, however, does not contain long IDR stretches (Fig. 1a). The limited linking regions of GIT1 also lack features of IDRs that have propensities to undergo phase separation^18^. Therefore, it is unlikely that GIT1 phase separation is driven by IDRs. Since the coiled-coil domain of GIT1 mediates its dimerization^37^ (Fig. 3f), we first investigated whether this interaction is required for GIT1 phase separation. A triple mutant of GIT1 (L438P, L459P, L466P; referred to as the ‘‘LP’’ mutant hereafter) converted the dimeric GIT1-CC into a monomer (Fig. 3f). The GIT1_LP mutant completely lost its capacity to form condensed droplets *in vitro* and in HeLa cells (Fig. 3g and 3h), indicating that GIT1 dimer formation is essential for its phase separation. However, dimerization alone is likely not sufficient in supporting GIT1 phase separation, as multivalent inter-molecular interactions are known to be required for phase separation of biomolecular complexes^2,16^. Therefore, we searched for additional molecular interaction(s) between GIT1. We found that the N-terminal region of GIT1 (aa 1-371, GIT1-NTD corresponding to the GAS tandem) specifically binds to its C-terminal half (aa 371-end, GIT1-CTD) (Fig. 3i). Since the coiled-coil domain of GIT1 adopts a parallel conformation during its dimerization, thus the binding between NTD and CTD of GIT1 is likely inter-molecular in nature based on the topology of the protein conformation. Interestingly, addition of the β-Pix GBD peptide blocked the interaction between NTD and CTD of GIT1 (Fig. 3i), likely because the β-Pix binding site and the CTD binding site on GIT1 NTD overlap with each other. It is predicted that the β-Pix GBD peptide should be able to prevent GIT1 phase separation by blocking GIT1 oligomerization. Indeed, addition of the β-Pix GBD peptide abolished the phase separation of Cy3-GIT1 *in vitro*, whereas a mutant β-Pix GBD peptide (GBD_Y542D), which is deficient in GIT1-binding, had no impact on GIT1 phase separation (Fig. 3g). Taken together, we conclude that both coiled-coil-mediated dimerization and the interaction between GIT1 NTD and CTD contribute to the phase separation of GIT1 (Fig. 3j).

### Binding of β-Pix to GIT1 promotes phase separation of GIT1

Since β-Pix is a strong binding partner of GIT1 and β-Pix can form stable trimer via its coiled-coil domain^37^, formation of the GIT1 and β-Pix complex would further increase GIT1 oligomerization thus might promote the phase separation of GIT1. Indeed, mixing equal molar amount of Cy3-labeled GIT1 with Alexa488-labeled β-Pix led to formation of condensed liquid droplets enriched with both proteins (Fig. 4a). Importantly, addition of β-Pix lowered the threshold concentration for GIT1 to undergo phase separation and dramatically increased the number of condensed droplets of GIT1 (Fig. 3a *vs* 4a). β-Pix alone at a concentration up to 100 μM did not undergo phase separation (data not shown). When GFP-GIT1 and RFP-β-Pix were co-expressed in HeLa cells, we observed many bright and completely overlapping spherical puncta enriched with both proteins (Fig. 4b). No puncta were observed in cells expressing RFP-β-Pix alone (Fig. 4b), indicating that β-Pix by itself could not form condensed phase. FRAP analysis showed that the GFP-GIT1 signal within the puncta could be recovered after photo bleaching, but only to approximately 20% of its original intensity within a few minutes (Fig. 4c). Notably, the exchange rate of GIT1 between the condensed phase and the dilute cytoplasmic phase was considerably slower than those reported in other phase separation systems in cells^7,10,38^, suggesting that the GIT1/β-Pix condensates are less dynamic possibly due to the very tight binding between the two proteins. The β-Pix_Y542D mutant has a ∼5,000-fold reduction in binding to GIT1 (Supplementary Fig. 3a). Interestingly, when GFP-GIT1 was co-expressed with β-Pix_Y542D, the recovery speed of GFP-GIT1 signal after photo-bleaching is much faster than that of GIT1 co-expressed with WT β-Pix (Fig. 4c and Supplementary Fig. 3b). Y542 of β-Pix was reported to be phosphorylated in cells by FAK or Src^39,40^. It is tempting to speculate that phosphorylation of β-Pix at Y542 may be a regulatory switch for the GIT1/β-Pix condensates to disperse, a hypothesis that needs to be tested in the future.

**Fig. 4.**
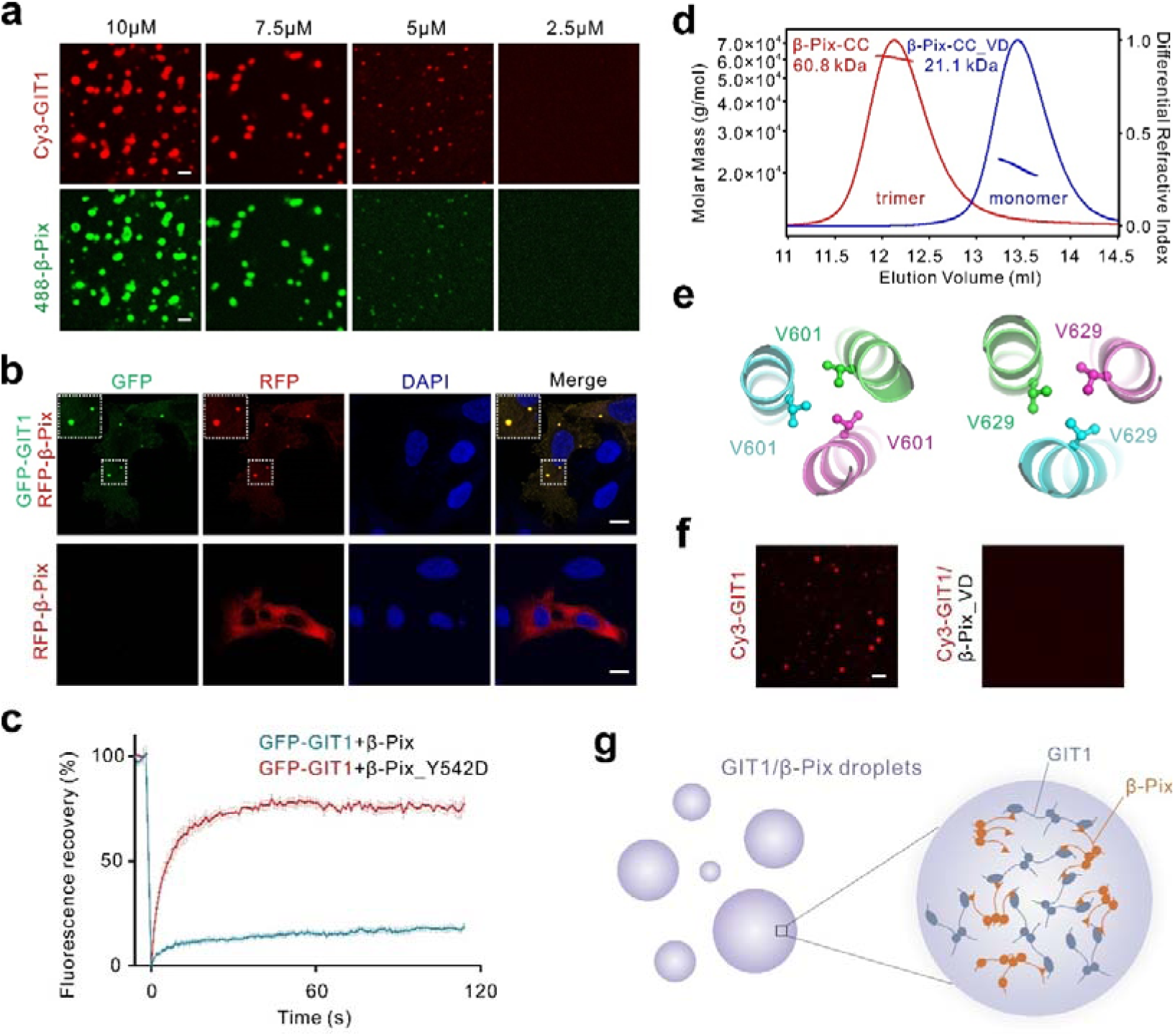
Formation of the GIT1/β-Pix condensates. (**a**) Fluorescence images showing that mixture of GIT1 and β-Pix (both proteins are in their full-length forms) led to phase separation at indicated concentrations. GIT1 and β-Pix were labeled with Cy3 and Alexa488 at 1%, respectively. Scale bar: 5 μm. (**b**) Representative images showing that co-expression of GFP-GIT1 and RFP-β-Pix in HeLa cells produced multiple spherical puncta, whereas RFP-β-Pix alone was diffused in cytoplasm. Scale bar: 5 μm. (**c**) FRAP analysis showing that GFP-GIT1 in the puncta, when co-expressed with WT RFP-β-Pix, exchanged slowly with the protein in dilute cytoplasm. The signal recover is also very limited. In contrast, the exchange of GFP-GIT1 between the puncta and cytoplasm, when co-expressed with Y542D mutant of β-Pix, was much faster and the signal recovery was also much higher. (**d**) FPLC-coupled static light-scattering analysis showing that wild type β-Pix-CC forms a stable trimer in solution, and the ‘‘VD’’ mutant is a monomer. (**e**) Close-up view of the interactions between V601 and V629 in β-Pix trimer (PDB code: 2W6B), respectively. (**f**) Fluorescence images showing that β-Pix_VD mutant abolished the phase separation of GIT1. The assay was under the same condition as in Fig. 3a with each protein at a concentration of 20 μM. (**g**) A schematic model showing the interaction network formed by GIT1 and β-Pix in the condensates. Scale bar: 5 μm.

We next investigated the role of β-Pix valency in promoting phase separation of the GIT1/β-Pix complex. The coiled-coil domain of β-Pix is a trimer in solution^37^. Guided by the structure (PDB code: 2W6B), we designed a two-point mutant of β-Pix-CC (i.e., V601D, V629D; referred to as the ‘‘VD’’ mutant hereafter) capable of converting trimeric β-Pix-CC into a monomer (Fig. 4d and 4e). Mixing Alexa488-labeled monomeric β-Pix_VD with Cy3-GIT1 did not promote phase separation of GIT1; instead, it eliminated phase separation of GIT1 (Fig. 4f). It is likely that the monomeric β-Pix_VD, analogous to the β-Pix GBD peptide (Fig. 3h and 3i), can specifically bind to GIT1 and consequently disrupt the NTD/CTD binding-mediated oligomerization of GIT1. The above data indicate that the phase separation of the GIT1/β-Pix complex is driven by formation of large molecular network contributed by the coiled-coil domain-mediated multimerization of both GIT1 and β-Pix as well as the very strong interaction between GIT1 and β-Pix (Fig. 4g).

### Paxillin’s connection with the GIT1/β-Pix complex

The best studied role for the GIT/PIX complexes is that in FAs and cell migrations. GIT1 is recruited to FAs via direct binding to Paxillin^41,42^. Paxillin is a multi-domain scaffold protein composed of five leucine-rich sequences known as LD motifs and four LIM (Lin11, Isl-1 and Mec-3) domains (Fig. 5a). The FAT domain of GIT1 has been reported to bind to LD2 and LD4 of Paxillin^43,44^. We confirmed that GIT1 FAT bind to LD2 and LD4 motifs with a *K*_d_ of ∼164 μM and ∼3.0 μM, respectively (Fig. 5a and Supplementary Fig. 4). The remaining three Paxillin LD motifs had no detectable binding to GIT1 FAT (Fig. 5a and Supplementary Fig. 4).

**Fig. 5.**
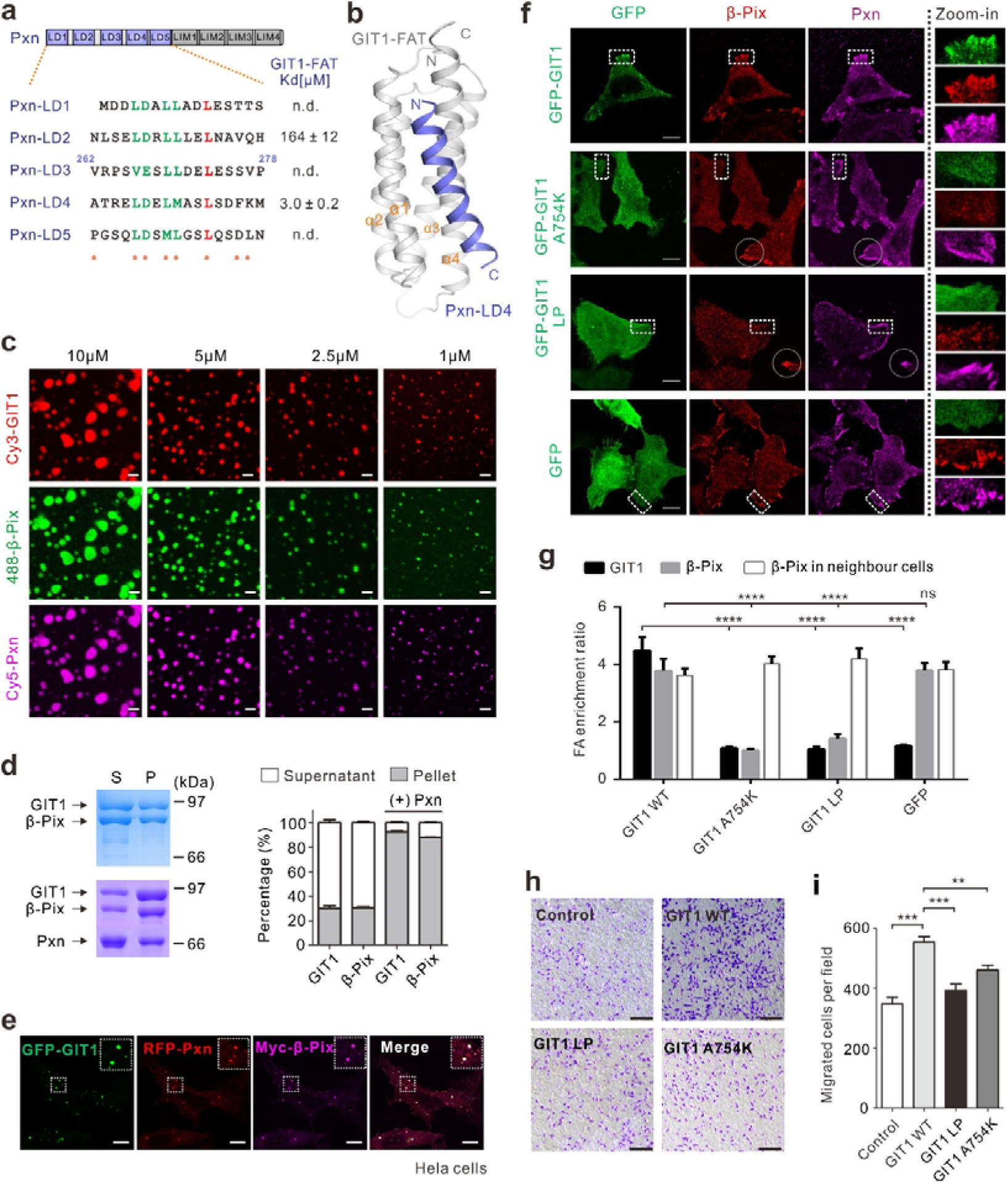
Paxillin further promotes GIT1/β-Pix phase separation. (**a**) Schematic diagram showing the domain organization of Paxillin. The sequence alignment of the LD motifs of Paxillin is included. The identical and conserved residues are color in red and green, respectively. Residues involved in the GIT1/Paxillin interaction are indicated by orange dots. The dissociation constants of the interactions between various Paxillin LD motifs and GIT1 FAT obtained from ITC-based assays were shown. (**b**) Ribbon diagram representation of the GIT1 FAT/Paxillin LD4 complex structure. (**c**) Fluorescence images showing that mixing Paxillin, GIT1 and β-Pix at indicated concentrations resulted in condensed droplets with three proteins simultaneously enriched in the condensed phase. Paxillin, GIT1 and β-Pix were labeled with Cy5, Cy3 and Alexa488, respectively, with each at 1% level. Scale bar: 5 μm. (**d**) Representative SDS-PAGE analysis and quantification data showing the distributions of GIT1 and β-Pix in the supernatant (S) and pellet (P) with or without presence of Paxillin in sedimentation-based assays. The final concentration of each protein was 5 μM. Results are expressed as mean ± SD from three independent batches of sedimentation experiments. (**e**) Representative images showing co-expression of GFP-GIT1, RFP-Paxillin and Myc-β-Pix in HeLa cells produced many spherical puncta with all three proteins co-localized together. Scale bar: 5 μm. (**f**) GFP-GIT1 could recruit endogenous β-Pix to FAs marked by anti-Paxillin antibody in HeLa cells. The GFP-GIT1^A754K^ or GFP-GIT1^LP^ mutants impaired the FA localization of β-Pix. The neighboring non-transfected cells worked as internal controls. The GFP only served as the vector control of the experiment. Scale bar: 5 μm. (**g**) Quantification of FA enrichments of GFP-GIT1 and its mutants as well as β-Pix derived from experiment described in (**f**). For each group, fifteen cells (i.e., n=15) from three independent batches were imaged for quantifications. The FA enrichment ratio is defined as [GFP_FA intensity_]/[GFP_cytoplasm intensity_] or [β-Pix_FA intensity_]/[β-Pix_cytoplasm intensity_], respectively, and is expressed as mean ± SEM for each group; ns, not significant; ****p < 0.0001, using one-way ANOVA with Dunnett’s multiple comparison test. (**h**) Transwell migration assays were performed to measure the cell migration activities of HeLa cells transfected with GFP-GIT1, GFP-GIT1^A754K^, GFP-GIT1_LP and GFP control. (**i**) Quantification of cell migration activities of GFP-GIT1 and its mutants from experiment described in (**h**). Data are expressed as mean ± SEM for each group from six independent experiments. ***p < 0.001, **p < 0.01, using one-way ANOVA with Dunnett’s multiple comparison test.

We determined the crystal structure of GIT1 FAT in complex with Paxillin LD4 (Table S1). In the complex, GIT1 FAT adopts a stable four-helix bundle structure, which is similar to the apo-form FAT structure (Fig. 5b and Supplementary Fig. 5a). The LD4 motif forms an α-helix occupying the binding site formed by α1 and α4 of GIT1 FAT (Fig. 5b). The GIT1 FAT/Paxillin LD4 interface is mainly mediated by hydrophobic interactions (see Supplementary Fig. 5b and 5c for detailed interactions). The GIT1 FAT/Paxillin LD4 interface is similar to those in other FAT/LD interactions such as the Pyk2/Paxillin, FAK/Paxillin and CCM3/Paxillin complexes (Supplementary Fig. 5d-f). Determination of the GIT1 FAT/Paxillin LD4 complex allowed us to design specific point mutations on either GIT1 or Paxillin leading to complete disruption of the complex formation (e.g., L669K^GIT1^, A754K^GIT1^, and F276D^Paxillin^, Supplementary Fig. 5g). These mutations were used to investigate the role of Paxillin in targeting the GIT1/β-Pix condensates to FAs and the role of the GIT1/β-Pix condensates in regulating cell migration (see below).

### Paxillin further promotes phase separation of the GIT1/β-Pix complex

Since both of the LD2 and LD4 motifs are able to bind to GIT1, binding of Paxillin can further expand the valency of the GIT/PIX complexes and thus promote its phase separation. Indeed, addition of the full-length Paxillin to the GIT1/β-Pix complex further lowered the threshold concentration for the GIT1/β-Pix complex to undergo phase separation and Paxillin was also recruited to the condensed phase of the GIT1/β-Pix complex (Fig. 5c). Formation of the condensed droplets was readily observed at an individual protein concentration of 1 μM or lower (Fig. 5c), suggesting that the Paxillin/GIT1/β-Pix condensates can form at their physiological concentrations. Consistent with the imaging-based analysis, the amount of GIT1 and β-Pix proteins in the condensed phase (the ‘‘pellet’’ fraction) significantly increased when Paxillin was added in a sedimentation-based assay^10^ (from ∼30% to ∼90%, Fig. 5d). To examine whether such co-puncta of three proteins may occur in living cells, GFP-GIT1, RFP-Paxillin and Myc-β-Pix were co-expressed in HeLa cells. Under fluorescence microscope, these three proteins formed many co-localized, micrometer-sized spherical puncta in cells (Fig. 5e). Taken together, the above studies demonstrated that the interaction between GIT1 and Paxillin further promotes phase separation of the GIT1/β-Pix complex.

### GIT1/β-Pix condensates are recruited to focal adhesions and required for cell migrations

Endogenous β-Pix was found to be enriched in FAs in cells with or without overexpression of GFP-tagged WT GIT1 (Fig. 5f, *top row*; quantified in Fig. 5g)^44^. In sharp contrast, overexpression of a Paxillin binding-deficient mutant of GIT1, GFP-GIT1^A754K^, led to dramatic decreased FA localization of the endogenous β-Pix (*second row* of Fig. 5f and Fig. 5g). As internal controls, the endogenous β-Pix could be effectively targeted to FAs in neighboring cells without expression of GIT1^A754K^ (Fig. 5f, highlighted with circles). Similarly, in the cells expressing GFP alone, the endogenous β-Pix co-clustered with Paxillin at FAs (Fig. 5f and 5g). The reduction of β-Pix FA localization in GFP-GIT1^A754K^ expressing cells is likely due to the dominant negative effect of the mutant GIT1, as the mutant is still believed to bind to β-Pix with a nanomolar dissociation constant. The above results indicate that the GIT1/β-Pix condensates formed by the endogenous level of both enzymes can be targeted to FAs by GIT1-mediated binding to Paxillin.

A critical question is whether formation of the clustered GIT1/β-Pix FA puncta in cells requires phase separation of the GIT1/β-Pix. To address this question, we took the advantage of the monomeric GIT1_LP mutant characterized in Fig. 3f-h. The GIT1_LP mutant is incapable of undergoing phase separation (Fig. 3g and 3h), but the mutant does not affect its binding to either β-Pix or Paxillin (Fig. 1 and Fig. 5a). Thus, the GIT1_LP mutant can be used to specifically assess the role of GIT1/β-Pix phase separation in the FA targeting of the complex. Satisfyingly, the endogenous β-Pix could not be recruited to FAs effectively when cells were expressing the GFP-GIT1_LP mutant (*third row* of Fig. 5f and Fig. 5g). Considering that regulation of FA dynamics is crucial for cell motility, one would expect that perturbation of recruitment of the GIT1/β-Pix condensates to FAs would impair cell migrations. Indeed, using a trans-well migration assay, we found that expression of GIT1 WT significantly promoted cell migrations. In contrast, neither GFP-GIT1^A754K^ nor the GFP-GIT1_LP mutant could promote cell migrations (Fig. 5h and 5i). The above data suggest that both the formation of the GIT1/β-Pix condensates via phase separation and Paxillin-mediated targeting of the condensates to FAs play a role in regulating cell motilities.

### GIT/PIX condensates regulate neuronal synapse formation via PIX C-terminal PDZ binding motif-mediated synaptic targeting

In addition to FAs, the GIT/PIX complexes are found in other cellular locations such as intracellular vesicles, neuronal synapses, centrioles, cell-cell junctions, DNA damage repair foci etc., where they regulate diverse cellular functions^23,45^. Since there are several protein-protein binding domains/motifs in both proteins, we hypothesized that GIT and PIX may use these domains/motifs to interact with various cellular proteins which can in turn target the GIT/PIX condensates to distinct cellular sites. For example, Scribble, a component of the Scribble/Lgl/Dlg master cell polarity regulatory complex^46^, can bind to and position β-Pix to specific subdomains in polarized cells^47,48^. Scribble contains a leucine-rich repeat (LRR) domain and four PDZ domains. The three N-terminal PDZ domains can bind to the PDZ-binding motif (PBM) of β-Pix with micromolar affinities^49^ (Fig. 6a). In neurons, the Shank family scaffold proteins can use their PDZ domains to bind to β-Pix and thus concentrate β-Pix to postsynaptic densities (PSDs) of excitatory synapses for Rac-dependent dendritic spine dynamic modulations^50^ (Fig. 6a).

**Fig. 6.**
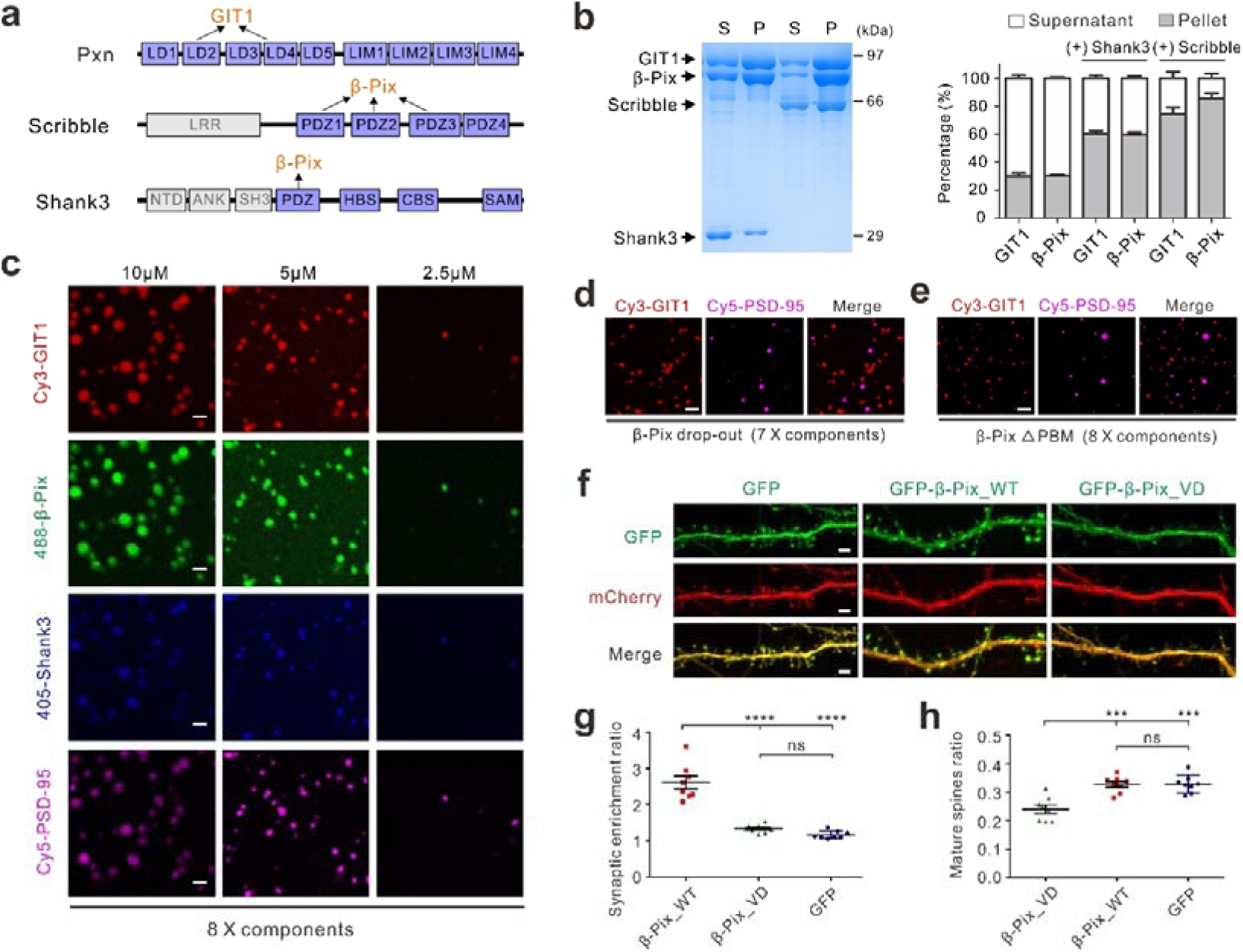
The GIT1/β-Pix condensate module can be recruited to synapses and required for dendritic spine development. (**a**) Schematic diagrams showing the domain organizations of Paxillin, Scribble, and Shank3. Domains that interact with GIT1 or β-Pix are indicated. (**b**) Sedimentation-based assays showing that Scribble and Shank3 could be enriched and in return promote phase separation of GIT1 and β-Pix. Results are expressed as mean ± SD from three independent batches of sedimentation experiments. (**c**) Fluorescence images showing that mixing GIT1, β-Pix and 6x PSD components (PSD-95, GKAP, Shank3, SynGAP, NR2B, and Homer) at indicated concentrations resulted in condensed droplets with eight proteins simultaneously enriched in the condensed phase. PSD-95, Shank3, GIT1 and β-Pix were labeled with Cy5, iFluor405, Cy3 and Alexa488, respectively, with each at 1% level. Scale bar: 5 μm. (**d**) Fluorescence images showing that GIT1 condensates did not overlap with the 6x PSD condensates when β-Pix was dropped out in the system. (**e**) Fluorescence images showing that GIT1/β-Pix^ΔPBM^ condensates did not overlap with the 6x PSD condensates. (**f**) Cultured hippocampal neurons were transfected with mCherry as the cell fill and GFP-tagged β-Pix constructs (GFP-β-Pix_WT, GFP-β-Pix_VD, GFP control) at DIV 14. After four days of expression, neurons were fixed and mounted for imaging. GFP-β-Pix_WT showed prominent spine localization, whereas the monomeric mutant β-Pix_VD had a diffused distribution pattern with no significant synaptic enrichment. Notably, compared to WT and GFP control group, neurons expressing β-Pix_VD mutant showed a severe reduction in the proportion of mature spines. (**g**) Quantification of the imaging data showing synaptic targeting of various β-Pix constructs. Synaptic enrichment ratio of β-Pix is defined as: [GFP_spine_/GFP_shaft_]/[mCherry_spine_/mCherry_shaft_]. Eight neurons (i.e., n=8) from three independent batches of cultures were imaged for each group, each neuron was analyzed for four branches (i.e., n=32). Error bar indicates ± SEM. ****p < 0.0001; ns, not significant. One-way ANOVA with Tukey’s multiple comparison test was used for the plot. (**h**) Quantification of image data showing reduction of mature spines for the neurons expressed the β-Pix_VD mutant. Eight neurons (i.e., n = 8) from three independent batches of cultures were imaged for each group for quantifications. Error bar indicates ± SEM. ***p < 0.001; ns, not significant. One-way ANOVA with Tukey’s multiple comparison test.

We used a sedimentation-based assay to test whether Scribble and Shank can be enriched in the GIT1/β-Pix condensates. We used purified Scribble PDZ1-4 for such assay, as the protein behaves well (e.g. highly soluble and non-aggregating, suitable for phase separation study). The Shank3 protein used in the assay contains PDZ-HBS-CBS-SAM as previously described^9^. For simplicity, we refer to these two proteins as Scribble and Shank3 here. When mixing either Scribble or Shank3 with GIT1 and β-Pix at a 1:1:1 molar ratio and each at 5 μM concentration, Scribble or Shank3 was readily recovered from the condensed phase (Fig. 6b), indicating that both proteins can be recruited and enriched into the GIT1/β-Pix condensates. Like what Paxillin does (Fig. 5d), both Scribble and Shank3 can further promote phase separation of the GIT1/β-Pix complex (Fig. 6b).

In a fluorescence imaging assay, iFluo405-Shank3 coalesced into the micrometer-sized GIT1/β-Pix condensates (Supplementary Fig. 6). Furthermore, the GIT1/β-Pix complex could be integrated into the excitatory PSD condensates consisting of PSD-95, GKAP, Shank3, SynGAP, NR2B, and Homer (i.e., the 6x PSD assembly in our earlier study, see ref 9) (Fig. 6c). Formation of the resulting 8-component condensates is specific, as addition of the LD4 peptide into the system did not affect phase separation of these synaptic proteins (Supplementary Fig. 7). Importantly, when β-Pix (the link between GIT1 and the PSD components) was dropped out in the 8x PSD mixture, GIT1 could still form condensates but the formed condensates no longer overlapped with the 6x PSD condensates (Fig. 6d). Moreover, the GIT1/β-Pix^ΔPBM^ condensates (β-Pix^ΔPBM^ binds to GIT1 as the WT β-Pix does, but the mutant cannot bind to Shank3) did not overlap with the 6x PSD condensates as well (Fig. 6e and Supplementary Fig. 8). The above results indicated that the specific Shank3/β-Pix interaction is required for recruitment of the GIT1/β-Pix condensates to the PSD condensates (i.e. Shank3 functions as the adaptor for targeting GIT1/β-Pix to the PSD condensates).

We next investigated whether phase separation of the GIT1/β-Pix complex is required for its synaptic targeting and function. In cultured hippocampal neurons, the expressed GFP-β-Pix_WT showed prominent spine localization, whereas the phase separation-deficient mutant of β-Pix, β-Pix_VD, which is a monomer but retains its bindings of β-Pix to GIT1 and Shank3 (Fig. 4d-f), had a diffused distribution pattern with no significant synaptic enrichment (Fig. 6f and quantified in Fig. 6g). Notably, compared to WT β-Pix and the GFP control, neurons expressing β-PIX_VD showed a severe reduction of mature spines (Fig. 6f and quantified in Fig. 6h), presumably due to disruption of the GIT/PIX condensates and subsequent defects in synaptic enrichment of this complex in neurons. Taken together, the above data indicate that the GIT1/β-Pix condensates are recruited to synapses likely via the β-Pix/Shank3 interaction and phase separation of the GIT1/β-Pix complex is crucial for synaptic targeting and dendritic spine development in hippocampal neurons.

## Discussion

In this study, we made two unexpected findings with potential general implications in cell biology. First, our results indicate that enzymes by themselves (in this case two small GTPase regulatory enzymes GIT1 and β-Pix) can form highly specific condensates via phase separation both *in vitro* and in living cells. Importantly, such condensates do not require additional scaffold proteins or scaffold-like molecules such as RNAs and DNAs, which are often essential for liquid-liquid phase separation in most of the currently known biological condensate systems^18,21,51^. Enzymes are generally extremely efficient bio-catalyzers and therefore do not need to exist at high concentrations inside the cell. Yet enzymes are often required to concentrate at targeted subcellular regions to perform location-specific cellular functions. Via liquid-liquid phase separation, enzymes such as GIT1 and β-Pix can autonomously form highly concentrated molecular assemblies, thus providing a novel mechanism for enriching limited amounts of enzymes into specific cellular regions for fast and spatially defined catalysis. Guided by the atomic structures of GIT1, β-Pix and the GIT1/β-Pix complex, we have also demonstrated that the phase separation-mediated GIT1/β-Pix complex condensation, instead of the classical binary interaction between GIT1 and β-Pix, is required for the enzyme complex to modulate cell migrations and synapse formation. It should be noted that formation of large enzyme complexes via classical stoichiometric interactions in cells (e.g. locally concentrated metabolic enzyme complexes^52^) has been known for decades (see ref 52 for a review). Our study provides a new paradigm to understand how such locally concentrated enzyme complexes might be formed via phase separation. Phase separation-mediated formation of dense enzyme complex condensates may have distinct properties in aspects such as enzyme kinetics, substrate accessibilities, threshold concentration and regulations of the enzyme assembly formation, etc. All these will need to be addressed in the future investigation.

Second, we show here that the GIT1/β-Pix condensates can function as a highly concentrated module capable of being recruited to diverse cellular signaling compartments by binding to specific adaptor proteins such as Paxillin, Scribble and Shank3 (Fig. 7). With this modular feature, the GIT1/β-Pix condensates can be specifically recruited by different adaptors to perform an array of cellular functions in various tissues or at different cell growth stages. The modular feature of the GIT1/β-Pix condensates for targeting the enzyme complex to different cellular processes is in a way analogous to the protein module-based cellular signal transduction pathway organizations. We suggest that formation of such modular regulatory enzyme condensates via phase separation may be a common mechanism for cells to utilize limited amounts of enzymes for broad and optimal cellular functions.

**Fig. 7.**
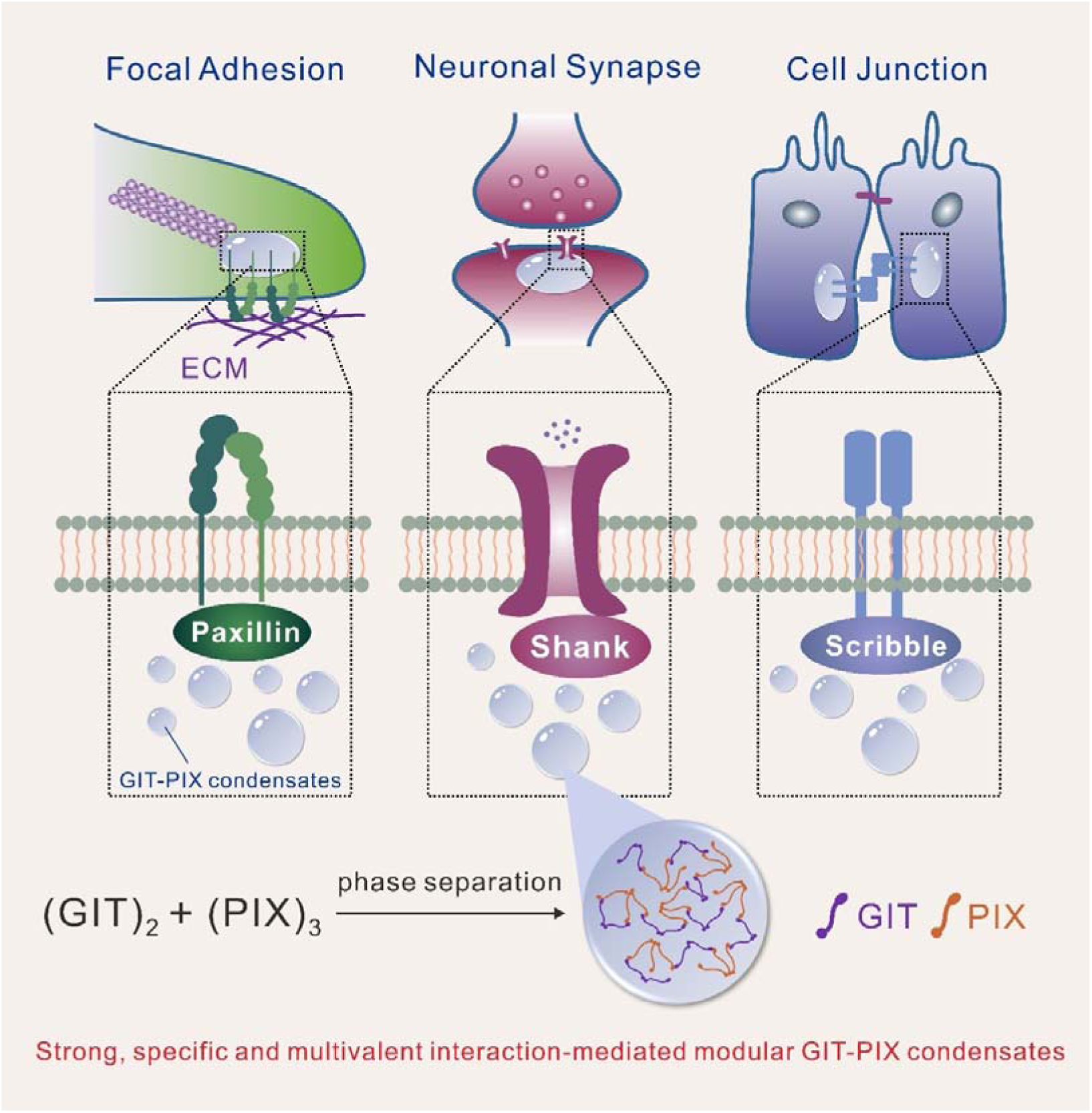
The versatile modular GIT/PIX condensates function in diverse cellular processes. A model depicting that the GIT/PIX condensates function as a modular organization capable of being targeted to distinct cellular compartments such as focal adhesions, neuronal synapses, and cell-cell junctions, enabling spatiotemporal regulation of GTPases activities. Note that the GIT/PIX condensates can be formed by strong, specific, and multivalent interactions between these two enzymes without additional scaffolding molecules.

In contrast to the majority of phase separation induced biomolecular condensates, the formation of the GIT1/β-Pix condensates requires very specific and strong interaction between these two proteins (*K*_d_ ∼15 nM). Such specific interaction is presumably in accordance with the specific functional roles of the two enzymes in various cellular processes. We argue that strong and specific multivalent interactions are critical for forming functionally specific biomolecular condensates in living cells, as we have demonstrated here and also in neuronal synapses previously^9,10^. Numerous studies in the past have illustrated the critical roles played by weak and multivalent interactions in the phase separation of biomolecules. Nonetheless, it is hard to envisage that formation of functionally specific biological signaling condensates is predominantly dictated by weak, promiscuous biomolecular interactions. It is also puzzling how genetic mutations that only lead to mild changes in the binding affinity of the corresponding protein, can cause human diseases when the interaction involved is weak and promiscuous. It is however possible that strong and specific interactions, together with weak but often high valency bindings, allow for the formation of highly specific biomolecular condensates. These assemblies may have very low phase separation concentration thresholds and broad dynamic properties. For this reason, we recommend the use of full-length proteins to investigate biomolecular condensates formation whenever possible. One thing is for certain, much remains to be uncovered in this exciting and emerging field of phase separation induced membrane-less biomolecular condensates.

Here, we have provided evidence that the GIT1/β-Pix condensates can be recruited to FAs via the GIT1/Paxillin interaction. Numerous observations point to the possibility that FAs, analogous to pre- and post-synaptic signaling assemblies in neurons^9,12^ and signaling complexes in immune-synapses^53^, may also be assembled via phase separation. Both cryo-electron tomography and recent super-resolution light microscopy studies have revealed that FAs form specific electron-dense structures. These condensed FA assemblies reside beneath the integrin-enriched plasma membrane and are open to the dilute cytoplasm^54–56^. Live-cell imaging studies have shown that the molecular assembly of FAs is extremely dynamic and fluid^56,57^. In particular, the actin cytoskeleton is intimately coupled to FA dynamics, at least in part via the GIT1/β-Pix complex-regulated GTPase signaling processes^34,45^. In the future, biochemical reconstitution will be a promising approach to directly test whether components of FAs can spontaneously assemble into biomolecular condensates.

## Acknowledgments

We thank the Shanghai Synchrotron Radiation Facility (SSRF, China) BL19U1 for X-ray beam time, the staff members of the Large-scale Protein Preparation System and Molecular Imaging System at the National Facility for Protein Science in Shanghai (NFPS), Zhangjiang Lab, China for providing technical support and assistance in data collection and analysis, Yuan Shang (University of Arizona, USA) for technical support during structure determination, Wenyu Wen (Fudan University, China) for providing assistance with the fluorescence polarization assay, Jinchuan Zhou for critical reading of the manuscript. This work was supported by grants from the National Key R&D Program of China (2018YFA0507900 to JZ and 2017YFA0504901 to RZ), a grant from National Natural Science Foundation of China (31770779) to JZ, a grant from the Chief Scientist Program of Shanghai Institutes for Biological Sciences, Chinese Academy of Sciences to RZ, and grants from RGC of Hong Kong (AoE-M09-12 and C6004-17G) and a grant from Asia Foundation for Cancer Research to MZ. MZ is a Kerry Holdings Professor in Science and a Senior Fellow of IAS at HKUST.

## Author contributions

JZ, QZ, and MZ designed the experiments. JZ, QZ, YX, and MP performed the experiments. YX, MP, LL, and JL carried out the X-ray data collection and structure determination. JZ, QZ, YX, LL, JL, RZ, and MZ analyzed the data. JZ, QZ and MZ drafted the manuscript. JZ and MZ coordinated the project. All authors approved the final version of the manuscript.

## METHODS

### Constructs and peptides

Mouse *GIT1* (GenBank: NM_001004144.1), *GIT2* (GenBank: NM_001077360.1), *Arhgef7* (encoding β-Pix; GenBank: NM_017402.4), *Pxn* (encoding Paxillin; GenBank: NM_133915.3), *Arf6* (GenBank: NM_007481.3) and *GGA1* (GenBank: NM_145929.2) genes were amplified from mouse brain cDNA library. Mouse full length *Shank3* gene was kindly provided by Dr. Guoping Feng at Massachusetts Institute of Technology. Human *Scrib* (encoding Scribble; GenBank: NM_015356.4) gene was amplified from human cDNA library. Various fragments of these genes were amplified by standard PCR method and cloned into pGEX 4T-1, pET32M3C (with a N-terminal Trx-His_6_ tag), pEGFP.N1, pTRFP or pCMV-Myc vector. Mutations were created through site-directed mutagenesis method. All constructs were confirmed by DNA sequencing.

The wild type β-Pix GBD peptide (sequence: ALEEDAQILKVIEAYCTSAKT), various β-Pix GBD mutant peptides, and LD4 peptide (sequence: ATRELDELMASLSDFKM) were commercially synthesized by ChinaPeptides (Shanghai, China) with purity > 95%.

### Protein expression and purification

Recombinant proteins were expressed in *Escherichia coli* BL21 (DE3) or (Codon Plus) cells at 16 °C for 18h inducing by the isopropyl-β-D-thiogalactoside (IPTG) at a final concentration of 0.2 mM. The N-terminal Trx-His_6_ tagged and GST-tagged proteins were purified by Ni^2+^-NTA agarose affinity chromatography and GSH-Sepharose affinity chromatography, respectively, and followed by a Superdex-200 26/60 size-exclusion chromatography. For β-Pix and Paxillin full length proteins, a step of monoQ column was used to remove nucleic acids contamination or degraded proteins.

### Isothermal titration calorimetry (ITC) assay

ITC measurements were carried out on a MicroCal VP-ITC system (Malvern) at 25 °C. Various GIT1 (in the cell, ∼50 μM) and β-Pix (in the syringe, ∼500 μM) proteins were in the buffer containing 50 mM Tris (pH 8.0), 100 mM NaCl, and 4 mM β-ME. LD motifs of Paxillin (in the syringe, ∼500 μM) and GIT1 FAT (in the cell, ∼50 μM) proteins were in the buffer containing 50 mM Tris, pH 8.0, 100 mM NaCl, 1 mM EDTA and 1 mM DTT. In each titration, 10 μL aliquot of protein in the syringe was injected into the cell at a time interval of 120 s make sure that the titration peak returned to the baseline. Titration data were fitted with the one-site binding model using Origin 7.0.

### GST-pull down assay

Flag-tagged wild type and mutants of GIT2 GAS were overexpressed in HEK293T cells. Cells were harvested and lysed by the ice-cold cell lysis buffer (50 mM HEPES pH 7.4, 150 mM NaCl, 10% glycerol, 2 mM MgCl_2_, 1% Triton and protease inhibitor cocktail). After centrifugation at 16,873 g for 10 min at 4 °C, the supernatants were incubated with 20 µl various wild type or mutants of GST-β-Pix GBD pre-loaded GSH-Sepharose 4B slurry beads. After extensive wash with the cell lysis buffer, the captured proteins were eluted by 20 µl 2× SDS-PAGE loading dye and detected by western blot using anti-Flag antibody (Sigma, 1:3000, Cat# F1804).

### Fast protein liquid chromatography (FPLC) coupled with static light scattering

The analysis was performed on an Agilent InfinityLab system coupled with a static light scattering detector (miniDawn, Wyatt) and a differential refractive index detector (Optilab, Wyatt). 150 μL GIT1-CC or β-Pix-CC protein sample at 50 μM was loaded into a Superose 12 10/300 GL column (GE Healthcare) pre-equilibrated with 50 mM Tris, pH 8.0, 100 mM NaCl, 1 mM EDTA, 1 mM DTT buffer. Data were analyzed using ASTRA 6 software (Wyatt).

### Fluorescence polarization assay

Fluorescence polarization assay was carried out on a PerkinElmer LS-55 fluorimeter equipped with an automated polarizer at 25 °C. In the assay, the commercially synthesized WT and mutant β-Pix GBD peptides were labeled with fluorescein-5-isothicyanate (FITC) (Invitrgen, Molecular Probe) at their N-termini. The FITC-labeled WT or mutant β-Pix GBD peptide was titrated with WT or mutant GIT2 GAS or GIT1 GAS in the buffer containing 50 mM Tris, pH 8.0, 100 mM NaCl, 1 mM DTT. The *K*_d_ value was fitted with classical one-site specific binding model using GraphPad Prism.

### Crystallization, Data collection and Structure determination

#### The GIT2 GAS/β-Pix GBD complex

For the GIT2 GAS^S255A/S256A^/β-Pix GBD complex, GIT2 GAS^S255A/S256A^ was mixed with a commercially synthesized β-Pix GBD peptide in a molar ratio of 1:1.3 (∼8 mg/ml) in the buffer of 50 mM Tris, pH 8.0, 100 mM NaCl, 2 mM DTT. The best crystals were obtained by the hanging drop diffusion method at 16 °C in the buffer condition containing 0.2 M NaF, 0.1 M Bis-Tris propane/citric acid pH 6.7 and 16% PEG3350. Crystals were soaked in crystallization solution containing 25% glycerol for cryo-protection. The diffraction data were collected at BL19U1 at Shanghai Synchrotron Radiation Facility (SSRF, China). The diffraction data were processed with the HKL3000 package^58^. The complex structure was solved by the molecular replacement method by PHASER^59^ using the structure of GAP-ANK tandem of ACAP1 (PDB code: 3JUE) as the searching model. Further refinement was performed using PHENIX^60^ and Coot^61^. The final refinement statistics of the complex structures were listed in Table S1.

#### The GIT1 FAT/Paxillin LD4 complex

To obtain stable GIT1/Paxillin complex, GIT1 FAT (aa 640-770) was fused with a “GSGSGSGSGS” linker and Paxillin LD4 (aa 261-282). The best crystals of the fusion protein (∼20 mg/ml) were obtained by the hanging drop diffusion method at 16 °C in the buffer containing 0.2 M (NH4)_2_SO_4_, 30% PEG4000. Before X-ray diffraction experiments, crystals were soaked in crystallization solution containing 25% glycerol for cryo-protection. The diffraction data were collected at BL19U1 at Shanghai Synchrotron Radiation Facility (SSRF, China), and processed with the HKL3000 package. Using the structure of the GIT1 FAT apo form structure (PDB code: 2JX0) as the search model, the initial structural model was solved using the molecular replacement method using the software suits of PHASER. Refinements were carried out using PHENIX. The dataset was twinned with a twin fraction of 0.37 as indicated by phenix.xtriage^60^. Twin refinement restraints were applied during the refinement. Coot was used for Paxillin peptide modeling and model adjustments. The final refinement statistics of the complex structures were listed in Table S1. All structural diagrams were prepared by PyMOL.

### Protein labeling with fluorophore

Purified proteins were exchanged into the NaHCO_3_ buffer containing 300 mM NaCl, 100 mM NaHCO_3_, pH 8.3, 4 mM β-ME using a HiTrap desalting column and concentrated to 5 mg/ml before reaction. Cy3/Cy5 NHS ester (AAT Bioquest) and Alexa 488 NHS ester (Thermo Fisher) were dissolved in DMSO and incubated with the corresponding protein at room temperate for 1h. The fluorophore was mixed with protein solution in 1:1 molar ratio. The labeling reaction was quenched by the 200 mM Tris, pH 8.2 buffer, and the labeled protein was separated with a HiTrap desalting column into buffer containing 50 mM Tris, pH 8.0, 300 mM NaCl, and 4 mM β-ME. Fluorescence labeling efficiency was measured by Nanodrop 2000 (Thermo Fisher).

### In vitro phase transition assay

All purified proteins were exchanged into the buffer containing 50 mM Tris, pH 8.0, 300 mM NaCl, and 4 mM β-ME. After centrifugation at 16,873 g for 10 min at 4 °C, samples were placed on ice prior to the phase transition assay.

For sedimentation-based assays, GIT1 protein, GIT1/β-Pix mixture, or GIT1/β-Pix/Paxillin mixture each with total volume of 50 μL was incubated at room temperature for 10 min. Then, the mixture was centrifuged at 16,873 g for 3 min at 22 °C. Samples from supernatant fraction and pellet fraction were analyzed by SDS-PAGE with Coomassie blue staining. Each assay was performed three times. The intensity of each band on SDS-PAGE was quantified by ImageJ and data were presented as mean ± SD.

For microscope-based assays, each sample was injected into a home-made chamber as described previously^10^ for DIC (Nikon eclipse 80i) or fluorescent imaging (Zeiss LSM 880).

### Fluorescence recovery after photo-bleaching assay

FRAP assay was performed on a Zeiss LSM 880 confocal microscope with a 40X oil objective. For *in vitro* FRAP experiments on fluorophore labeled proteins, Cy3 signal was bleached by 561 nm laser beam at room temperature. For FRAP assay on puncta in living cell, HeLa cells were cultured on glass-bottom dishes (MatTek) and transfected with the indicated plasmids. GFP signal was bleached with 488 nm laser beam at 37 °C.

For each experiment, the fluorescence intensities of a neighboring droplet with similar size to the bleached one were also recorded for intensity correction. Background was subtracted before data analysis. The ROI intensity at time 0s (right after the photobleaching) was set as 0% and the pre-bleaching intensity was normalized to 100%.

### HeLa cell imaging, FA localization and cell migration

HeLa cells were cultured on 12-well plates and transfected with the indicated plasmids using Viafect (Promega, Madison, WI). After expression for 24h, cells were fixed with 4% paraformaldehyde (PFA) and immunostained with the indicated antibodies. Images were acquired on Leica SP8 or Zeiss LSM 880 confocal microscope by a 40× oil lens. Images were processed and analyzed using ImageJ. For FA localization analysis, three independent experiments were conducted in a blinded fashion. Focal adhesion regions were outlined and selected based on the Paxillin channel. The FA enrichment ratio was calculated as [GFP_FA intensity_]/[GFP_cytoplasm intensity_] or [β-Pix_FA intensity_]/[β-Pix_cytoplasm intensity_], respectively.

Cell migration experiment was performed using Transwell membrane filter inserts (8 μm pore size, Corning costar). 1 × 10^5^ HeLa cells were seeded into the upper chamber and allowed to migrate into the lower chamber for 16-18h at 37°C. Cells in the upper chamber were carefully wiped by cotton buds, cells at the bottom of the membrane were washed once with PBS, and fixed by 100% methanol for 10 min, and then stained with Crystal Violet Staining Solution (Beyotime Biotechnology) for 10 min. The migrated cells were counted under a light microscope from five random fields of each well. All experiments were performed three times.

### Primary hippocampal neuron culture and imaging

Hippocampal neuron cultures were prepared as previously described^62^. At DIV14, neurons were transfected with 2 mg indicated plasmids per well (12-well plate) using Lipofectamine 2000 reagent (Invitrogen). Neurons were fixed at DIV18 with 4% paraformaldehyde (PFA) together with 4% sucrose in 1 × PBS buffer and mounted on slides for imaging. Confocal images were obtained using a Leica SP8 confocal microscope with a 40 × oil-immersion lens. Transfected neurons were chosen randomly for quantification from at least three independent batches of cultures. For detailed spine visualization, an additional 4 × zoom factor was applied. Normally, four randomly selected dendrites (∼65 μm in length each) were imaged and analyzed from an individual neuron. Each image was collected as a z series maximum projection with 0.35-μm depth intervals. Intensity was measured with ImageJ.

### Quantification and statistical analysis

Statistical parameters including the definitions and exact values of n (e.g., number of experiments), distributions and deviations are reported in the Figures and corresponding Figure Legends. For FA localization and transwell migration assay, the results were expressed as mean ± SEM; ns, not significant, ****p < 0.0001, ***p < 0.001, **p < 0.01, using one-way ANOVA with Dunnett’s multiple comparison test. For synaptic targeting and spine development assay, the results were expressed as mean ± SEM; ns, not significant, ***p < 0.001, ****p < 0.0001, using one-way ANOVA with Tukey’s multiple comparison test. Statistical analysis was performed by GraphPad Prism.

### Data availability

The atomic coordinates of the GIT2/β-Pix and GIT1/Paxillin complexes are deposited to the Protein Data Bank under the accession codes: 6JMT and 6JMU, respectively. Other data are available from the corresponding author upon reasonable request.

## Supplemental Information

**Fig. S1:**
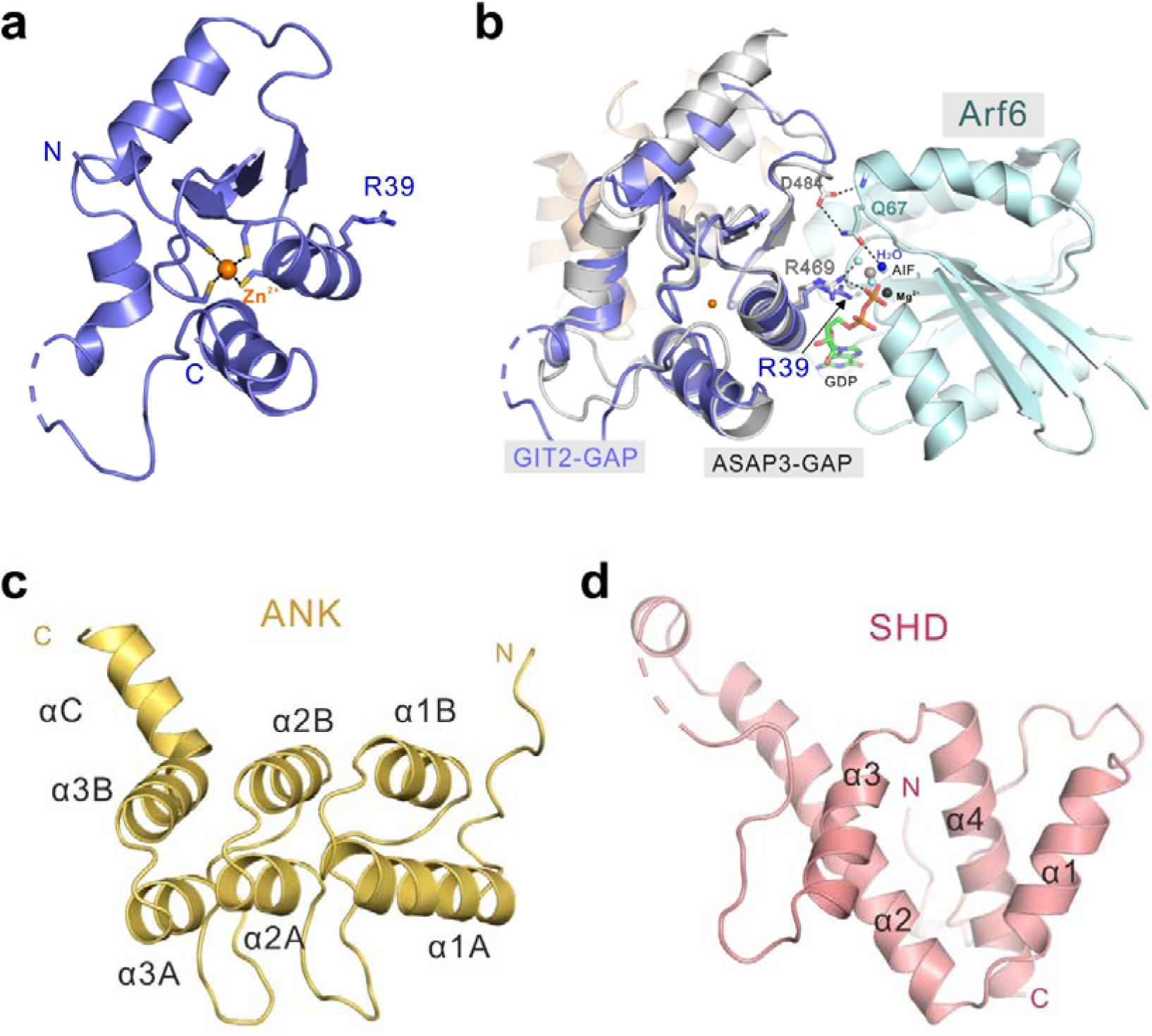
Structural analysis of ArfGAP, ANK, and SHD domains of GIT2. **a,** Ribbon diagram representation of the structure of ArfGAP domain. A zinc ion coordinated by four cystines is shown. The arginine finger, R39, is shown in stick mode. **b,** Superimposition of the structure of GAP^GIT2^ with that of GAP^ASAP3^ (PDB code: 3LVQ) showing that a conserved arginine of GAP^GIT2^, R39^GIT2^, aligns well with the arginine finger of ASAP3, R469^ASAP3^, which is required for GTP hydrolysis. **c,d,** Ribbon diagrams representation of the structures of ANK (**c**) and SHD (**d**) domains.

**Fig. S2:**
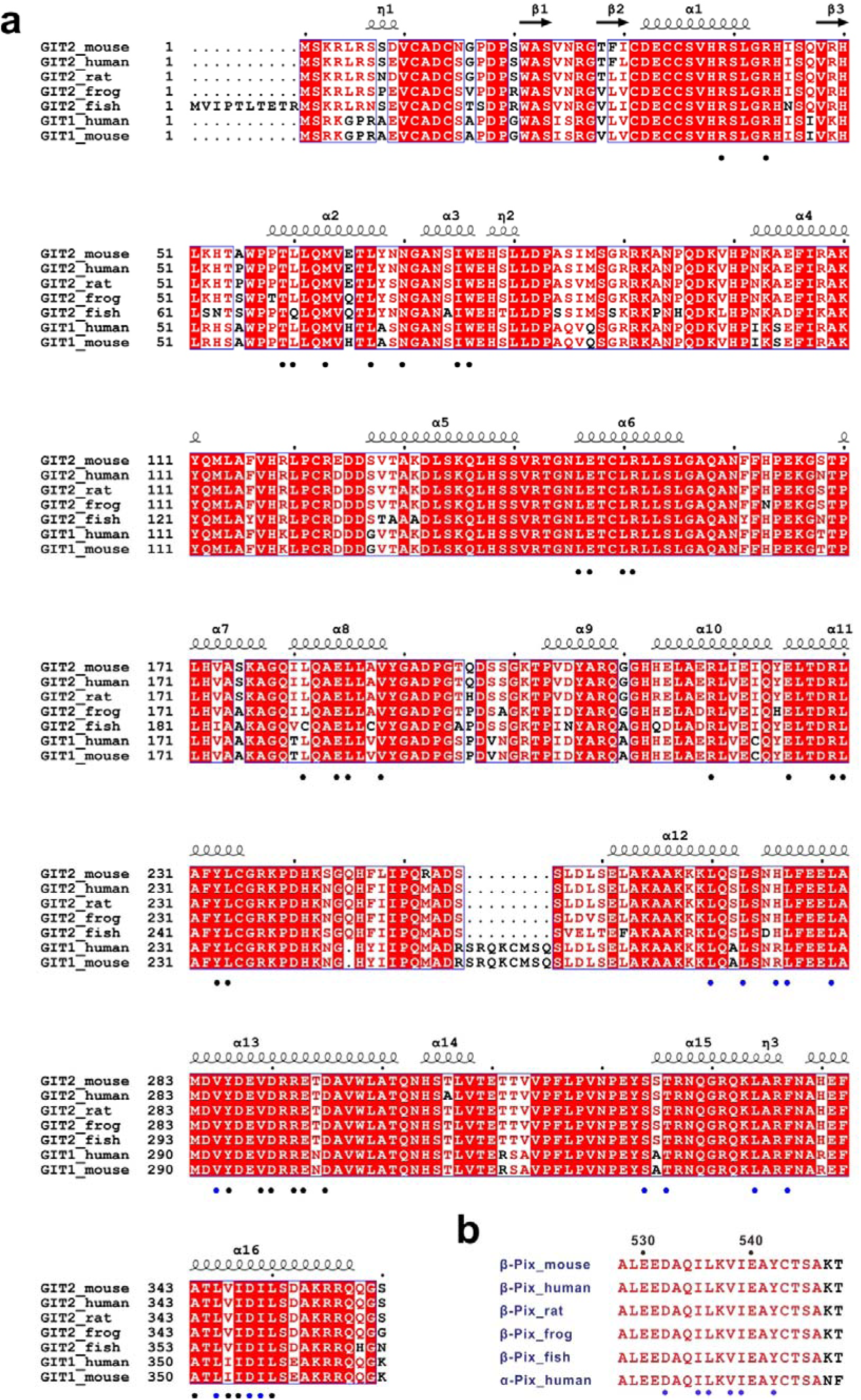
Sequence analysis of GIT and PIX family proteins. **a,** Structure-based sequence alignment of GIT family proteins. In this alignment, the identical residues are highlighted with red boxes, and the conserved residues are color in red. Residues required for domain-domain coupling in GIT GAS tandem and PIX-GBD binding are indicated by black and blue dots, respectively. **b,** Sequence alignment of GBD domains of PIX family proteins. The identical residues are color in red. Residues required for GIT GAS binding are indicated by blue dots.

**Fig. S3:**
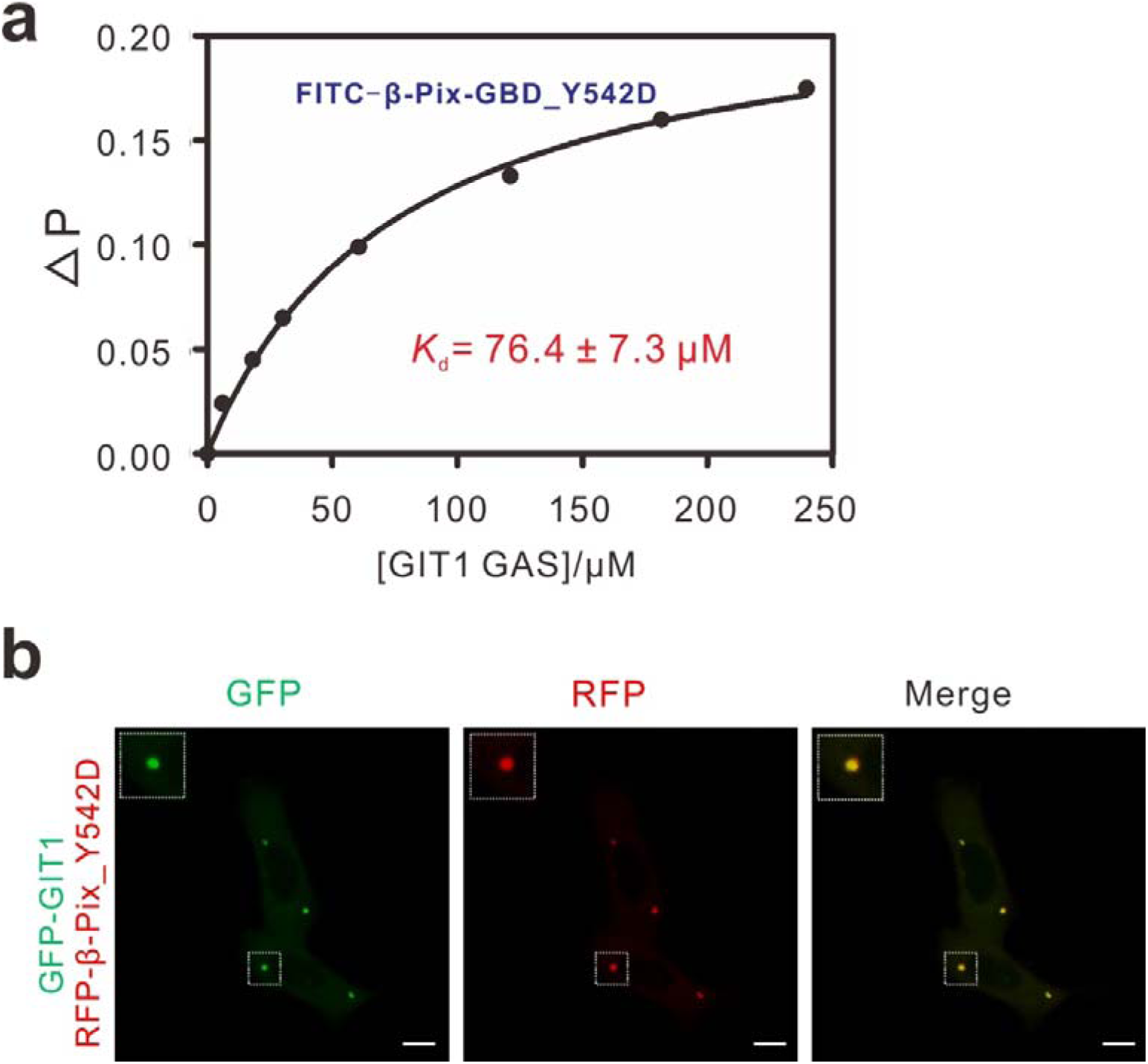
The Y542D mutation of β-Pix co-puncta with GIT1 in cells. **a,** A fluorescence polarization assay showing that the β-Pix-GBD_Y542D mutant bound with GIT1 GAS with a *K*_d_ of ∼76.4 μM. **b,** Representative images showing that co-expression of GFP-GIT1 and RFP-β-Pix_Y542D in HeLa cells produced multiple spherical puncta. Scale bar: 10 μm.

**Fig. S4:**
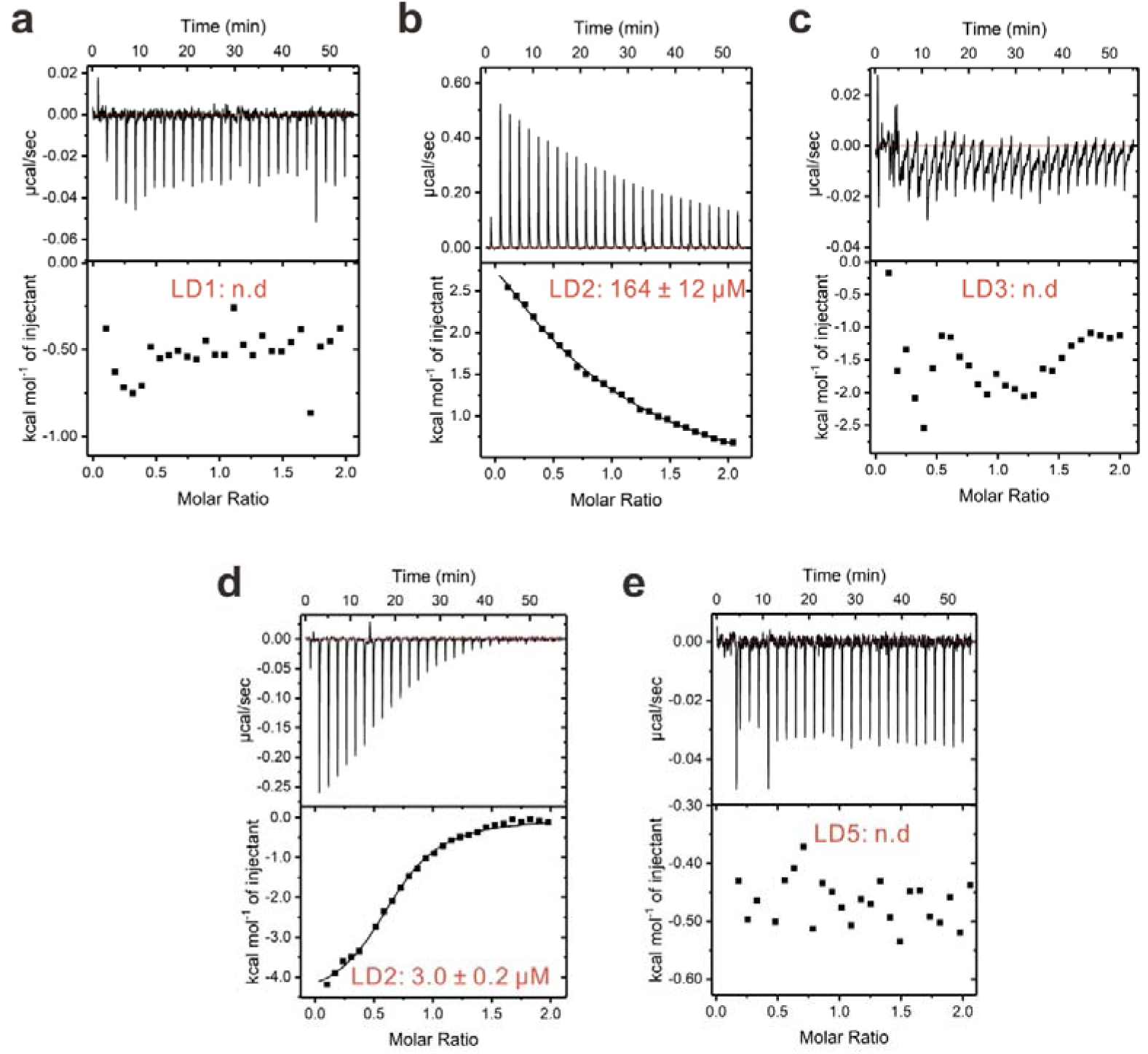
ITC titration curves showing the bindings of Paxillin LD motifs to GIT1-FAT. ITC-based binding curves of LD1 (a), LD2 (b), LD3 (c), LD4 (d), LD5 (e) to GIT1-FAT.

**Fig. S5:**
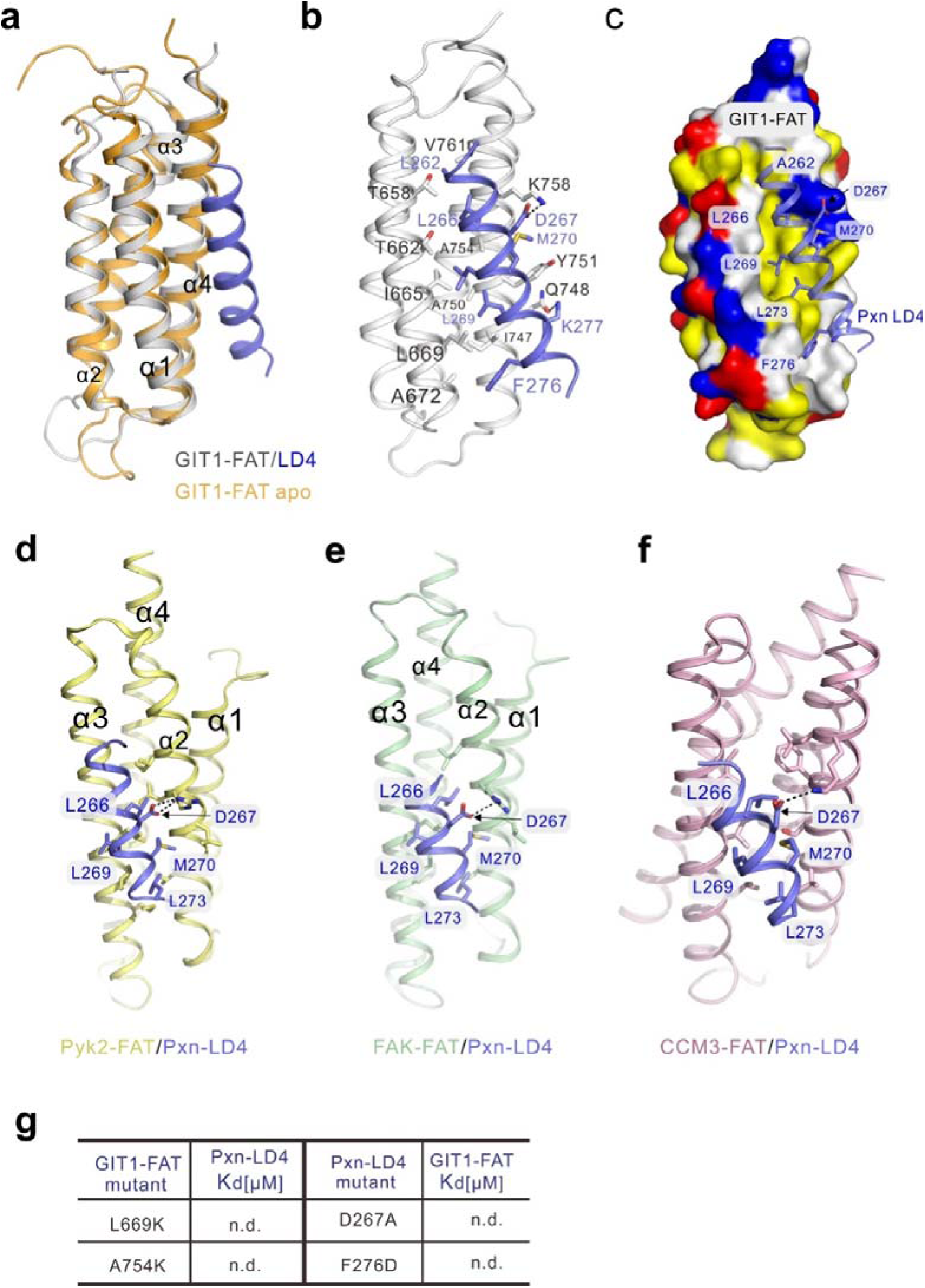
Structural analysis of the GIT1/Paxillin complex. **a,** Superimposition of the structures of the GIT1 FAT/Paxillin LD4 complex and apo GIT1 FAT. **b,** Detailed interactions between GIT1 FAT and Paxillin LD4. **c,** The combined surface and ribbon representations of the GIT1 FAT/Paxillin LD4 interface showing that the binding is mainly mediated by hydrophobic interactions. **d-f,** Structures of Pky2 FAT/LD4 (PDB code: 4R32) (**d**), FAK FAT/LD4 (PDB code: 1OW6) (**e**), CCM3 FAT/LD4 (PDB code: 3RQG) (**f**). **g,** ITC-based assays showing that mutations of selected key residues involved in the GIT1 FAT/Paxillin LD4 interface abolished the complex formation.

**Fig. S6:**
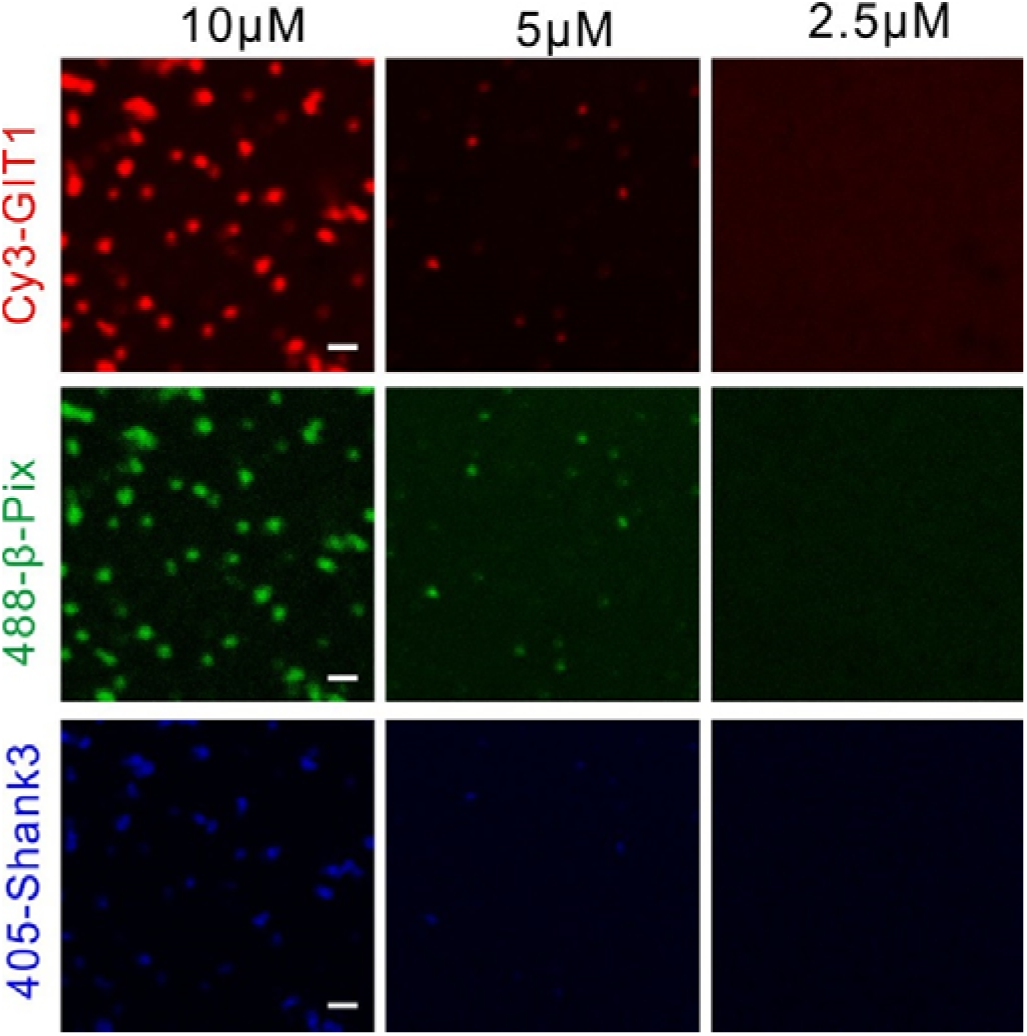
Shank3, GIT1, and β-Pix form condensed droplets. Fluorescence images showing that mixing GIT1, β-Pix and Shank3 at indicated concentrations resulted in condensed droplets with three proteins simultaneously enriched in the condensed phase. Shank3, GIT1 and β-Pix were labeled with iFluor405, Cy3 and Alexa488, respectively, with each at 1% level. Scale bar: 5 μm.

**Fig. S7:**
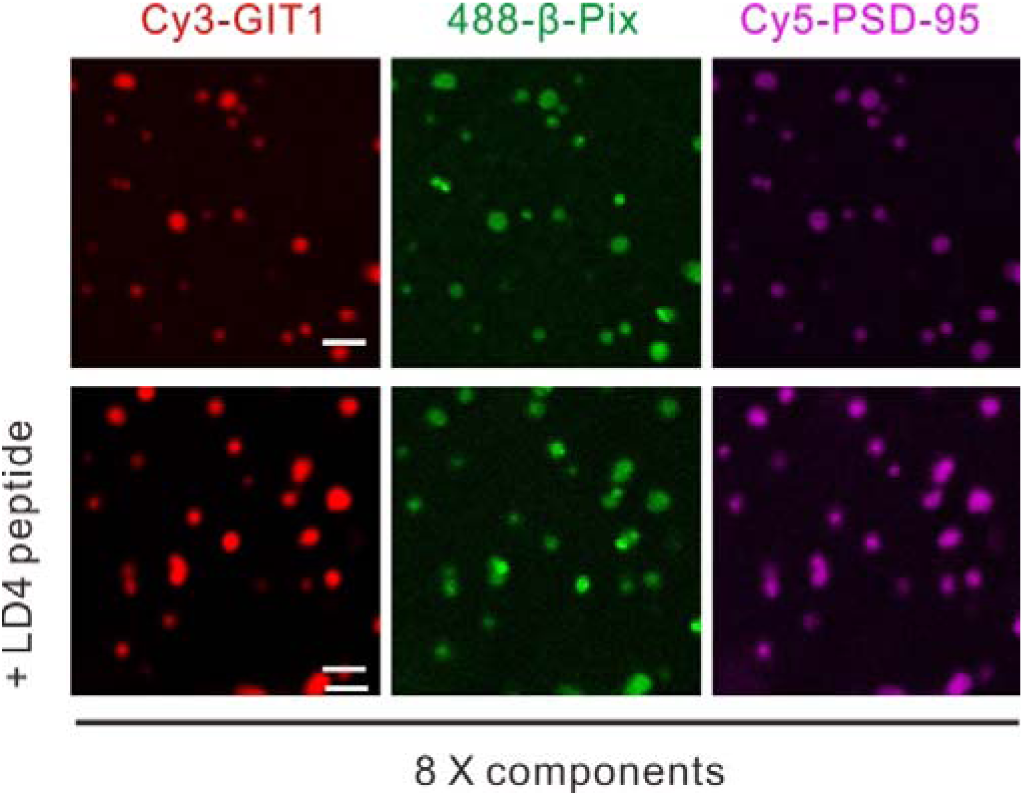
LD4 peptide did not affect the 8-component PSD condensate formation. Fluorescence images showing that addition of the LD4 peptide (200 μM) into the mixture of GIT1 and β-Pix (both proteins are in their full-length forms and each at the concentration of 5 μM) did not affect the phase separation of the two proteins. GIT1 and β-Pix were labeled with Cy3 and Alexa488 at 1%, respectively. Scale bar: 10 μm.

**Fig. S8:**
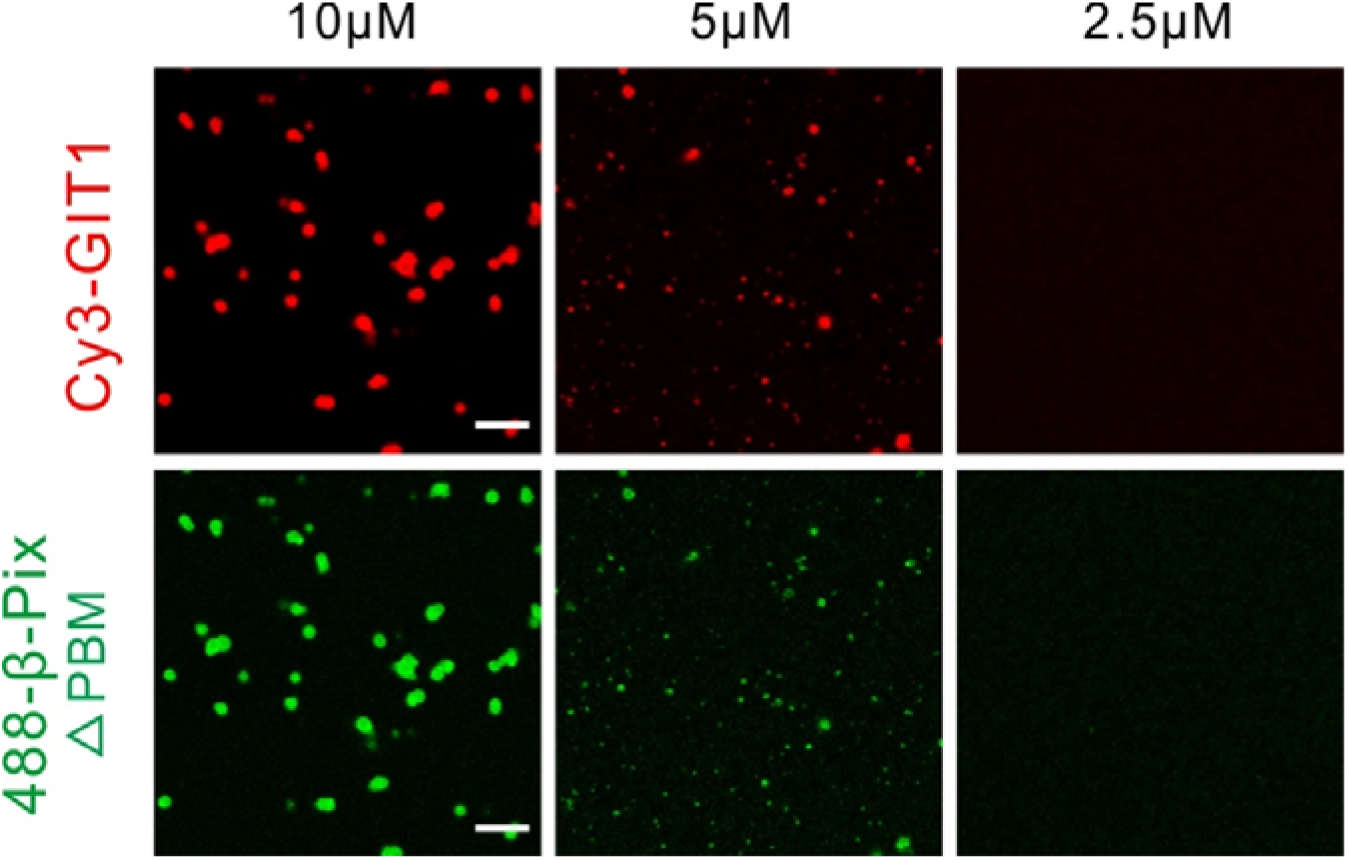
Formation of the GIT1/β-Pix^ΔPBM^ condensates. Fluorescence images showing that mixture of GIT1 and β-Pix^ΔPBM^ led to phase separation at indicated concentrations. GIT1 and β-Pix^ΔPBM^ were labeled with Cy3 and Alexa488 at 1%, respectively. Scale bar: 10 μm.

**Table S1.**
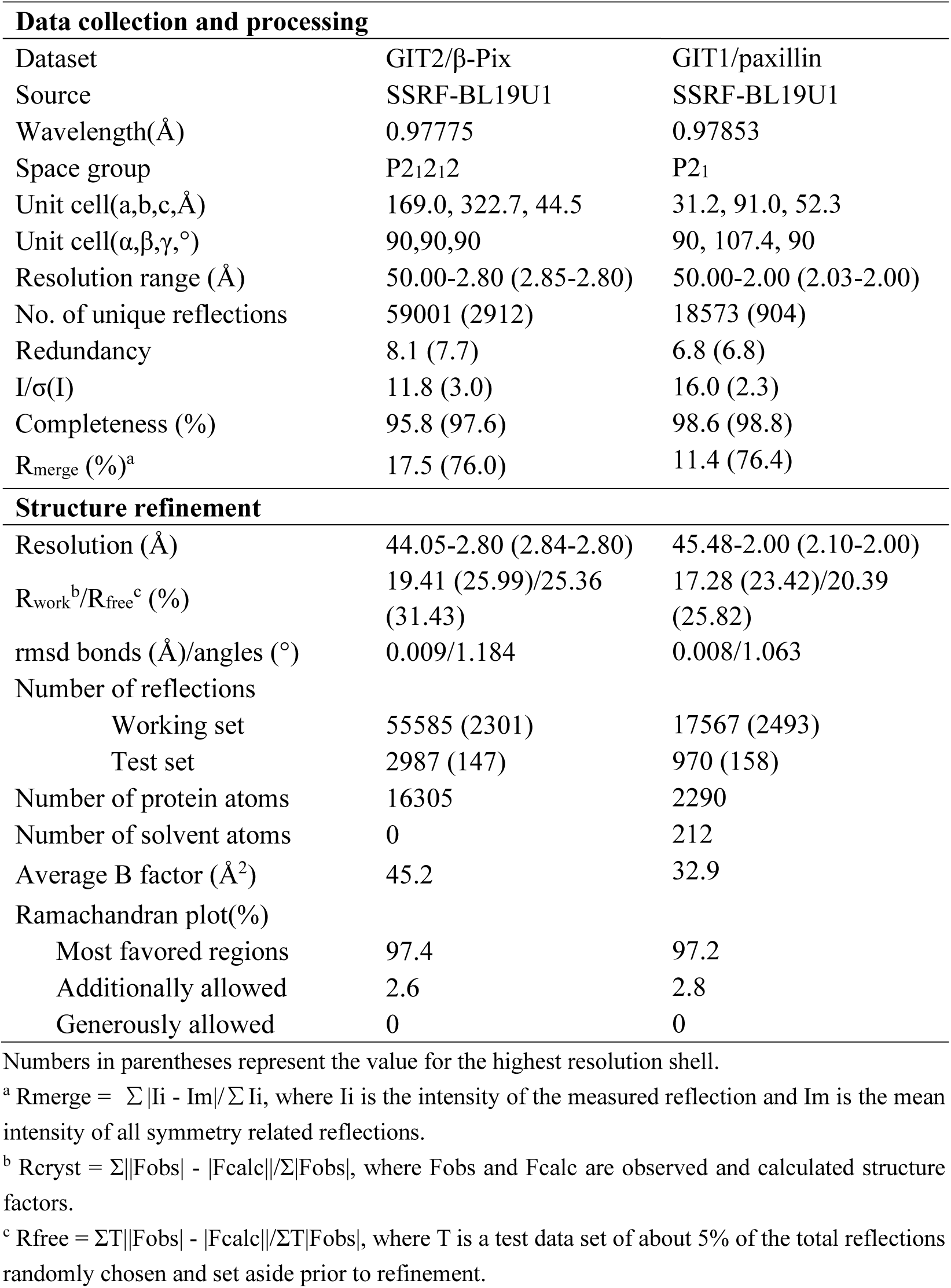
Data collection and refinement statistics.

| **Data collection and processing** |  |  |
| --- | --- | --- |
| Dataset | GIT2/β-Pix | GIT1/paxillin |
| Source | SSRF-BL19U1 | SSRF-BL19U1 |
| Wavelength($\text{\AA}$) | 0.97775 | 0.97853 |
| Space group | P2<sub>1</sub>2<sub>1</sub>2 | P2<sub>1</sub> |
| Unit cell(a,b,c,$\text{\AA}$) | 169.0, 322.7, 44.5 | 31.2, 91.0, 52.3 |
| Unit cell($\alpha,\beta,\gamma$,$^\circ$) | 90,90,90 | 90, 107.4, 90 |
| Resolution range ($\text{\AA}$) | 50.00-2.80 (2.85-2.80) | 50.00-2.00 (2.03-2.00) |
| No. of unique reflections | 59001 (2912) | 18573 (904) |
| Redundancy | 8.1 (7.7) | 6.8 (6.8) |
| I/$\sigma$(I) | 11.8 (3.0) | 16.0 (2.3) |
| Completeness (%) | 95.8 (97.6) | 98.6 (98.8) |
| R<sub>merge</sub> (%)<sup>a</sup> | 17.5 (76.0) | 11.4 (76.4) |
| **Structure refinement** |  |  |
| Resolution ($\text{\AA}$) | 44.05-2.80 (2.84-2.80) | 45.48-2.00 (2.10-2.00) |
| R<sub>work</sub><sup>b</sup>/R<sub>free</sub><sup>c</sup> (%) | 19.41 (25.99)/25.36<br>(31.43) | 17.28 (23.42)/20.39<br>(25.82) |
| rmsd bonds ($\text{\AA}$)/angles ($^\circ$) | 0.009/1.184 | 0.008/1.063 |
| Number of reflections |  |  |
| Working set | 55585 (2301) | 17567 (2493) |
| Test set | 2987 (147) | 970 (158) |
| Number of protein atoms | 16305 | 2290 |
| Number of solvent atoms | 0 | 212 |
| Average B factor ($\text{\AA}^2$) | 45.2 | 32.9 |
| Ramachandran plot(%) |  |  |
| Most favored regions | 97.4 | 97.2 |
| Additionally allowed | 2.6 | 2.8 |
| Generously allowed | 0 | 0 |

Numbers in parentheses represent the value for the highest resolution shell.
<sup>a</sup> R<sub>merge</sub> = $\sum |I_i - \bar{I}| / \sum I_i$ , where $I_i$ is the intensity of the measured reflection and $\bar{I}$ is the mean intensity of all symmetry related reflections.
<sup>b</sup> R<sub>cryst</sub> = $\sum ||F_{obs}| - |F_{calc}|| / \sum |F_{obs}|$ , where $F_{obs}$ and $F_{calc}$ are observed and calculated structure factors.
<sup>c</sup> R<sub>free</sub> = $\sum T ||F_{obs}| - |F_{calc}|| / \sum T |F_{obs}|$ , where T is a test data set of about 5% of the total reflections randomly chosen and set aside prior to refinement.

## References

1. Hyman, A.A., Weber, C.A. & Julicher, F. Liquid-liquid phase separation in biology. Annu Rev Cell Dev Biol 30, 39–58 (2014).

2. Banani, S.F., Lee, H.O., Hyman, A.A. & Rosen, M.K. Biomolecular condensates: organizers of cellular biochemistry. Nat Rev Mol Cell Biol 18, 285–298 (2017).

3. Shin, Y. & Brangwynne, C.P. Liquid phase condensation in cell physiology and disease. Science 357(2017).

4. Feng, Z., Chen, X., Wu, X. & Zhang, M. Formation of biological condensates via phase separation: Characteristics, analytical methods, and physiological implications. J Biol Chem (2019).

5. Brangwynne, C.P. et al. Germline P granules are liquid droplets that localize by controlled dissolution/condensation. Science 324, 1729–32 (2009).

6. Brangwynne, C.P., Mitchison, T.J. & Hyman, A.A. Active liquid-like behavior of nucleoli determines their size and shape in Xenopus laevis oocytes. Proc Natl Acad Sci U S A 108, 4334–9 (2011).

7. Woodruff, J.B. et al. The Centrosome Is a Selective Condensate that Nucleates Microtubules by Concentrating Tubulin. Cell 169, 1066–1077 e10 (2017).

8. Woodruff, J.B., et al. Centrosomes. Regulated assembly of a supramolecular centrosome scaffold in vitro. Science 348, 808–12 (2015).

9. Zeng, M. et al. Reconstituted Postsynaptic Density as a Molecular Platform for Understanding Synapse Formation and Plasticity. Cell 174, 1172–1187 e16 (2018).

10. Zeng, M. et al. Phase Transition in Postsynaptic Densities Underlies Formation of Synaptic Complexes and Synaptic Plasticity. Cell 166, 1163–1175 e12 (2016).

11. Feng, Z., Chen, X., Zeng, M. & Zhang, M. Phase separation as a mechanism for assembling dynamic postsynaptic density signalling complexes. Curr Opin Neurobiol 57, 1–8 (2018).

12. Wu, X. et al. RIM and RIM-BP Form Presynaptic Active-Zone-like Condensates via Phase Separation. Mol Cell (2019).

13. Patel, A. et al. A Liquid-to-Solid Phase Transition of the ALS Protein FUS Accelerated by Disease Mutation. Cell 162, 1066–77 (2015).

14. Molliex, A. et al. Phase separation by low complexity domains promotes stress granule assembly and drives pathological fibrillization. Cell 163, 123–33 (2015).

15. Boeynaems, S. et al. Protein Phase Separation: A New Phase in Cell Biology. Trends Cell Biol 28, 420–435 (2018).

16. Li, P. et al. Phase transitions in the assembly of multivalent signalling proteins. Nature 483, 336–40 (2012).

17. Banjade, S. & Rosen, M.K. Phase transitions of multivalent proteins can promote clustering of membrane receptors. Elife 3(2014).

18. Ditlev, J.A., Case, L.B. & Rosen, M.K. Who’s In and Who’s Out-Compositional Control of Biomolecular Condensates. J Mol Biol 430, 4666–4684 (2018).

19. Wang, J. et al. A Molecular Grammar Governing the Driving Forces for Phase Separation of Prion-like RNA Binding Proteins. Cell 174, 688–699 e16 (2018).

20. Chong, S. et al. Imaging dynamic and selective low-complexity domain interactions that control gene transcription. Science 361(2018).

21. Jain, A. & Vale, R.D. RNA phase transitions in repeat expansion disorders. Nature 546, 243–247 (2017).

22. Shan, Z. et al. Basal condensation of Numb and Pon complex via phase transition during Drosophila neuroblast asymmetric division. Nat Commun 9, 737 (2018).

23. Zhou, W., Li, X. & Premont, R.T. Expanding functions of GIT Arf GTPase-activating proteins, PIX Rho guanine nucleotide exchange factors and GIT-PIX complexes. J Cell Sci 129, 1963–74 (2016).

24. Premont, R.T. et al. beta2-Adrenergic receptor regulation by GIT1, a G protein-coupled receptor kinase-associated ADP ribosylation factor GTPase-activating protein. Proc Natl Acad Sci U S A 95, 14082–7 (1998).

25. Manser, E. et al. PAK kinases are directly coupled to the PIX family of nucleotide exchange factors. Mol Cell 1, 183–92 (1998).

26. Premont, R.T. et al. The GIT/PIX complex: an oligomeric assembly of GIT family ARF GTPase-activating proteins and PIX family Rac1/Cdc42 guanine nucleotide exchange factors. Cell Signal 16, 1001–11 (2004).

27. Zhao, Z.S., Manser, E., Loo, T.H. & Lim, L. Coupling of PAK-interacting exchange factor PIX to GIT1 promotes focal complex disassembly. Mol Cell Biol 20, 6354–63 (2000).

28. Peng, H. et al. MAT2B-GIT1 interplay activates MEK1/ERK 1 and 2 to induce growth in human liver and colon cancer. Hepatology 57, 2299–313 (2013).

29. Kutsche, K. et al. Mutations in ARHGEF6, encoding a guanine nucleotide exchange factor for Rho GTPases, in patients with X-linked mental retardation. Nat Genet 26, 247–50 (2000).

30. Won, H. et al. GIT1 is associated with ADHD in humans and ADHD-like behaviors in mice. Nat Med 17, 566–72 (2011).

31. Chang, D. et al. Accounting for eXentricities: analysis of the X chromosome in GWAS reveals X-linked genes implicated in autoimmune diseases. PLoS One 9, e113684 (2014).

32. Ismail, S.A., Vetter, I.R., Sot, B. & Wittinghofer, A. The structure of an Arf-ArfGAP complex reveals a Ca2+ regulatory mechanism. Cell 141, 812–21 (2010).

33. Vitale, N., et al. GIT proteins, A novel family of phosphatidylinositol 3,4, 5-trisphosphate-stimulated GTPase-activating proteins for ARF6. J Biol Chem 275, 13901–6 (2000).

34. Hoefen, R.J. & Berk, B.C. The multifunctional GIT family of proteins. J Cell Sci 119, 1469–75 (2006).

35. Mandiyan, V., Andreev, J., Schlessinger, J. & Hubbard, S.R. Crystal structure of the ARF-GAP domain and ankyrin repeats of PYK2-associated protein beta. EMBO J 18, 6890–8 (1999).

36. Wright, P.E. & Dyson, H.J. Intrinsically disordered proteins in cellular signalling and regulation. Nat Rev Mol Cell Biol 16, 18–29 (2015).

37. Schlenker, O. & Rittinger, K. Structures of dimeric GIT1 and trimeric beta-PIX and implications for GIT-PIX complex assembly. J Mol Biol 386, 280–9 (2009).

38. Sabari, B.R. et al. Coactivator condensation at super-enhancers links phase separation and gene control. Science 361(2018).

39. Feng, Q. et al. Cool-1 functions as an essential regulatory node for EGF receptor- and Src-mediated cell growth. Nat Cell Biol 8, 945–56 (2006).

40. Mayhew, M.W. et al. Identification of phosphorylation sites in betaPIX and PAK1. J Cell Sci 120, 3911–8 (2007).

41. Turner, C.E. et al. Paxillin LD4 motif binds PAK and PIX through a novel 95-kD ankyrin repeat, ARF-GAP protein: A role in cytoskeletal remodeling. J Cell Biol 145, 851–63 (1999).

42. Premont, R.T., Claing, A., Vitale, N., Perry, S.J. & Lefkowitz, R.J. The GIT family of ADP-ribosylation factor GTPase-activating proteins. Functional diversity of GIT2 through alternative splicing. J Biol Chem 275, 22373–80 (2000).

43. Zhang, Z.M., Simmerman, J.A., Guibao, C.D. & Zheng, J.J. GIT1 paxillin-binding domain is a four-helix bundle, and it binds to both paxillin LD2 and LD4 motifs. J Biol Chem 283, 18685–93 (2008).

44. Schmalzigaug, R. et al. GIT1 utilizes a focal adhesion targeting-homology domain to bind paxillin. Cell Signal 19, 1733–44 (2007).

45. Frank, S.R. & Hansen, S.H. The PIX-GIT complex: a G protein signaling cassette in control of cell shape. Semin Cell Dev Biol 19, 234–44 (2008).

46. Bilder, D., Li, M. & Perrimon, N. Cooperative regulation of cell polarity and growth by Drosophila tumor suppressors. Science 289, 113–6 (2000).

47. Audebert, S. et al. Mammalian Scribble forms a tight complex with the betaPIX exchange factor. Curr Biol 14, 987–95 (2004).

48. Dow, L.E. et al. The tumour-suppressor Scribble dictates cell polarity during directed epithelial migration: regulation of Rho GTPase recruitment to the leading edge. Oncogene 26, 2272–82 (2007).

49. Lim, K.Y.B., Godde, N.J., Humbert, P.O. & Kvansakul, M. Structural basis for the differential interaction of Scribble PDZ domains with the guanine nucleotide exchange factor beta-PIX. J Biol Chem 292, 20425–20436 (2017).

50. Park, E. et al. The Shank family of postsynaptic density proteins interacts with and promotes synaptic accumulation of the beta PIX guanine nucleotide exchange factor for Rac1 and Cdc42. J Biol Chem 278, 19220–9 (2003).

51. Du, M. & Chen, Z.J. DNA-induced liquid phase condensation of cGAS activates innate immune signaling. Science 361, 704–709 (2018).

52. Srere, P.A. Complexes of sequential metabolic enzymes. Annu Rev Biochem 56, 89–124 (1987).

53. Su, X. et al. Phase separation of signaling molecules promotes T cell receptor signal transduction. Science 352, 595–9 (2016).

54. Patla, I. et al. Dissecting the molecular architecture of integrin adhesion sites by cryo-electron tomography. Nat Cell Biol 12, 909–15 (2010).

55. Kanchanawong, P. et al. Nanoscale architecture of integrin-based cell adhesions. Nature 468, 580–4 (2010).

56. Winograd-Katz, S.E., Fassler, R., Geiger, B. & Legate, K.R. The integrin adhesome: from genes and proteins to human disease. Nat Rev Mol Cell Biol 15, 273–88 (2014).

57. Zaidel-Bar, R. & Geiger, B. The switchable integrin adhesome. J Cell Sci 123, 1385–8 (2010).

58. Minor, W., Cymborowski, M., Otwinowski, Z. & Chruszcz, M. HKL-3000: the integration of data reduction and structure solution--from diffraction images to an initial model in minutes. Acta Crystallogr D Biol Crystallogr 62, 859–66 (2006).

59. McCoy, A.J., et al. Phaser crystallographic software. J Appl Crystallogr 40, 658–674 (2007).

60. Adams, P.D. et al. PHENIX: a comprehensive Python-based system for macromolecular structure solution. Acta Crystallogr D Biol Crystallogr 66, 213–21 (2010).

61. Emsley, P. & Cowtan, K. Coot: model-building tools for molecular graphics. Acta Crystallogr D Biol Crystallogr 60, 2126–32 (2004).

62. Zhu, J. et al. Synaptic Targeting and Function of SAPAPs Mediated by Phosphorylation-Dependent Binding to PSD-95 MAGUKs. Cell Rep 21, 3781–3793 (2017).

